# Hyperactivated Glycolysis Drives Spatially-Patterned Kupffer Cell Depletion in MASLD

**DOI:** 10.1101/2025.09.26.678483

**Authors:** Jia He, Ran Li, Cheng Xie, Xiane Zhu, Keqin Wang, Zhao Shan

## Abstract

Metabolic dysfunction-associated steatotic liver disease (MASLD) progression is characterized by hepatic inflammation and cell death, yet the mechanisms underlying Kupffer cells (KCs) loss remain poorly understood. Here, we sought to elucidate the metabolic basis of KCs death during MASLD. Using metabolomics, immunostaining, and flow cytometry, we evaluated metabolic alterations and KCs death throughout early MASLD progression. We found that KCs death is an early hallmark of MASLD, exhibiting greater susceptibility and displaying a spatial distribution consistent with KCs zonation. Moreoever, KCs undergo progressive metabolic reprogramming toward enhanced glucose utilization during MASLD development, which is correlated with KCs death. In combination of biochemical agonist, isotope tracing and primary KCs culture, we further demonstrated that augmented glycolytic metabolism directly drives KCs death *in vitro*. Consistently, using *Chil1*-deficient mice, we further demonstrated that increased glucose utilization accelerates KCs *death in vivo*. Together, these findings establish a causal link between glycolytic activation and KCs loss during MASLD progression, highlighting glucose metabolic pathways as potential therapeutic targets to preserve KCs homeostasis and mitigate MASLD.

## Introduction

Metabolic dysfunction-associated steatotic liver disease (MASLD) emerges as a prevalent liver condition in the western world, affecting roughly one-third of the population^1–3^. Its incidence continues to climb due to its strong association with obesity, type 2 diabetes, and the metabolic syndrome^4–6^. MASLD includes a spectrum of liver disorders, from metabolic dysfunction-associated fatty liver (MAFL), often referred to as steatosis, to metabolic dysfunction-associated steatohepatitis (MASH)^7^. MASH, characterized by steatosis, inflammation, ballooning injury, and varying degrees of fibrosis, marks the initial critical stage of MASLD^8^. Kupffer cells (KCs) are liver-resident macrophages located in the hepatic sinusoids^9,10^. Derived from the embryonic yolk sac, KCs can self-renew through proliferation during adult homeostasis^11^. Studies suggest that KCs contribute to triglyceride storage during MASH progression, as demonstrated by depleting CD207^+^ KCs in CD207-DTR mice using diphtheria toxin (DT) or by using CD207^ΔBcl2l1^ mice to stimulate a low embryo-derived KCs (EmKCs) status^12^. Upon KCs death, monocyte-derived macrophages (MoMFs) gradually seed the KCs pool and eventually replace the deceased KCs^13^. These MoMFs tend to be more inflammatory than EmKCs, altering the liver’s response in MASH, eventually limiting triglyceride storage and contributing to liver fibrosis^12,14^. Researchers, including our team, have observed a gradual decline in KCs during MASLD development^12,14–17^. However, little is known regarding the dynamic loss of KCs and metabolic changes behind KCs death during MASLD.

Emerging evidence highlights a fundamental role of glucose metabolism in regulating macrophage function, polarization, and survival^18–21^. Metabolic reprogramming is a well-established hallmark of macrophage activation and functional adaptation across diverse contexts^22^. Within the unique metabolic milieu of the MASLD liver, characterized by lipotoxicity, insulin resistance, and disrupted nutrient fluxes, it is plausible that KCs metabolism is profoundly altered. However, whether these metabolic perturbations directly contribute to the observed loss of KCs, and through which specific metabolic pathways, remains unclear. Therefore, this study aims to investigate the role of glucose metabolism in governing KCs susceptibility to death during MASLD progression. We sought to define the glucose metabolic alterations that occur in KCs as MASLD develops and to establish a causal link between metabolic reprogramming and KCs loss. Elucidating these mechanisms is essential for advancing our understanding of MASLD pathogenesis and for identifying potential therapeutic targets to preserve KCs homeostasis and mitigate disease progression.

## Materials and methods

### Animal experiments and procedures

*Animals Chil1^-/-^* (strain no. T014402), *Chil1^flox//flox^* (strain no. T013652), and *Clec4f-cre* [C-type lectin domain family 4-Cre (*Clec4f-cre*), strain no. T036801] mice with a *C57BL/6J* background were purchased from GemPharmatech. Accordingly, *C57BL/6J* mice (strain no. N000013) were used as wild-type (WT) mice. To generate *Clec4f^△Chil1^* mice, *Chil1^flox//flox^* mice were crossed with *Clec4f-cre* mice and knock out efficiency was examined in KCs previously^17^. All mouse colonies were maintained at the Animal Core Facility of Yunnan University. The animal studies were approved by the Yunnan University Institutional Animal Care and Use Committee (IACUC, Approval No. YNU20220314). Male mice aged 6-8 weeks were used in this study. For 2-Deoxy-D-glucose (2-DG) Treatment in mice, WT C57BL/6J mice were fed a HFHC diet for five weeks. Beginning at week three, mice received intraperitoneal injections of either vehicle or 2-DG (50 mg/kg) every other day for two weeks. Mice were then euthanized for analysis of KCs viability and liver pathology.

*Construction of MASLD/MASH mouse model* Mice were provided a high-fat and high-cholesterol diet (HFHC, Research Diet, d12108c, 40 kcal% fat and 1.25% cholesterol) or a high-fat diet (HFD, Research Diet, d12492, 60 kcal% fat). Another group of mice was fed a methionine and choline deficient diet (MCD, Research Diet, A02082002BR). Throughout the feeding period, the body weight and food consumption of the mice were observed and recorded weekly. Once the dietary intervention was completed, the mice were euthanized. Liver and murine serum samples were collected for further analysis. Alanine aminotransferase (ALT) and aspartate aminotransferase (AST) levels in the serum, as well as cholesterol (TC) and triglyceride (TG) levels in both serum and liver tissues, were quantified using commercially available kits (Nanjing Jiancheng Bioengineering Institute). Genotyping Sample preparation and procedure were conducted as previously described^17^.

### Data Presentation and Statistical Analysis

Data in graph figures are presented as mean ± standard error of the mean (SEM). Statistical analyses were performed using SPSS Statistics (Version 22). For comparisons between two groups, an unpaired two-tailed Student’s t-test was used, while one-way analysis of variance (ANOVA) was applied for comparisons involving three or more groups. A p-value < 0.05 was considered statistically significant, with p-values indicated where applicable. All cell culture experiments were repeated at least three times independently. Figure 7 was created using the Figdraw platform (www.figdraw.com).

### Additional Methods

Additional detailed methods can be found in the Supporting Information.

## Results

### The death of Kupffer cells is a pathological characteristic during MASLD

To systematically investigate KCs death, we established an MASLD mouse model by feeding wildtype C57Bl/6J mice a high-fat high-cholesterol diet (HFHC) (Figure S1A). We then performed Hematoxylin and eosin (H&E), Oil Red O and Sirius red staining to assess immune cell infiltration, fat accumulation, and liver fibrosis in mice fed HFHC for 0, 4 or 16 weeks. After 4 weeks of HFHC feeding, we observed a slight increase in lipid droplets, with no apparent signs of liver inflammation or fibrosis in hepatocytes (Figure S1A). By 16 weeks of HFHC feeding, both lipid droplets and immune cell infiltration had increased significantly, though liver fibrosis --- as indicated by Sirius red staining, which labels collagen deposition --- was still not induced, and MASLD activity score remained below 4 (Figure S1A). Moreover, the body weight of mice fed HFHC gradually increased over the feeding period (Figure S1B). Analysis of serum alanine aminotransferase (ALT), aspartate aminotransferase (AST), serum and liver cholesterol or liver triglyceride (TG) levels revealed significant increases compared to mice fed a normal chow diet (NCD), except for serum TG, which was similar between NCD and HFHC-fed mice (Figure S1C). These findings collectively indicate the successful establishment of an early MASLD mouse model, without fibrosis development.

Subsequently, we investigated KCs death by labeling KCs with antibodies targeting T cell immunoglobulin mucin protein 4 (TIM4)^12,23^, and employing TdT-mediated dUTP Nick-End Labeling (TUNEL) to detect dead cells. Compared to baseline (0 weeks, prior to HFHC diet), KCs are significantly reduced with MASLD progression. Approximately 25% and nearly 60% of KCs underwent cell death by 4 and 16 weeks post-HFHC feeding, respectively (Figure 1A, 1B). To further validate KCs death during MASLD progression, we performed co-staining of the apoptotic marker cleaved caspase-3 (Cl-Casp3) with C-type lectin domain family 4 (Clec4f), another marker specific for KCs^13^. Although the ratio of Cl-Casp3^+^/Clec4f^+^ was relatively lower, its occurence and gradual increase followed a trend similar to that of the TUNEL staining (Figure S2A). Considering the potential recruitment of MoMFs into the liver, some of which may acquire KCs features, including TIM4 and Clec4f expression^12,14^, we performed flow cytometry analysis of liver nonparenchymal cells (NPCs) to distinguish resident KCs from monocyte-derived populations. Resident KCs were identified as CD45^+^ F4/80^hi^ CD11b^low^ TIM4^hi^ cells, MoKCs as CD45^+^ F4/80^hi^ CD11b^low^ TIM4^low^ cells, and infiltrating macrophages (IMs) as CD45^+^ F4/80^+^ CD11b^hi^ TIM4^-^ cells (Figure 1C). Our analysis showed that all TIM4^+^ KCs were of EmKCs, with no detectable MoKCs at any time point examined (Figure 1C). Consistent with the immunostaining results, the number of resident KCs decreased as early as 4 weeks and continued to decline through 16 weeks of HFHC feeding (Figure 1C, 1D). In contrast, IMs progressively accumulated during HFHC feeding (Figure 1C, 1D). To further refine the identification of MoMFs, we exclude CD45^+^ F4/80^+^ CD11b^hi^ TIM4^-^ Ly6G^+^ neutrophils from the IMs gate. This refined analysis yielded similar results, confirming a gradual increase in MoMFs over the course of HFHC feeding (Figure S2B, S2C).

**Figure 1.**
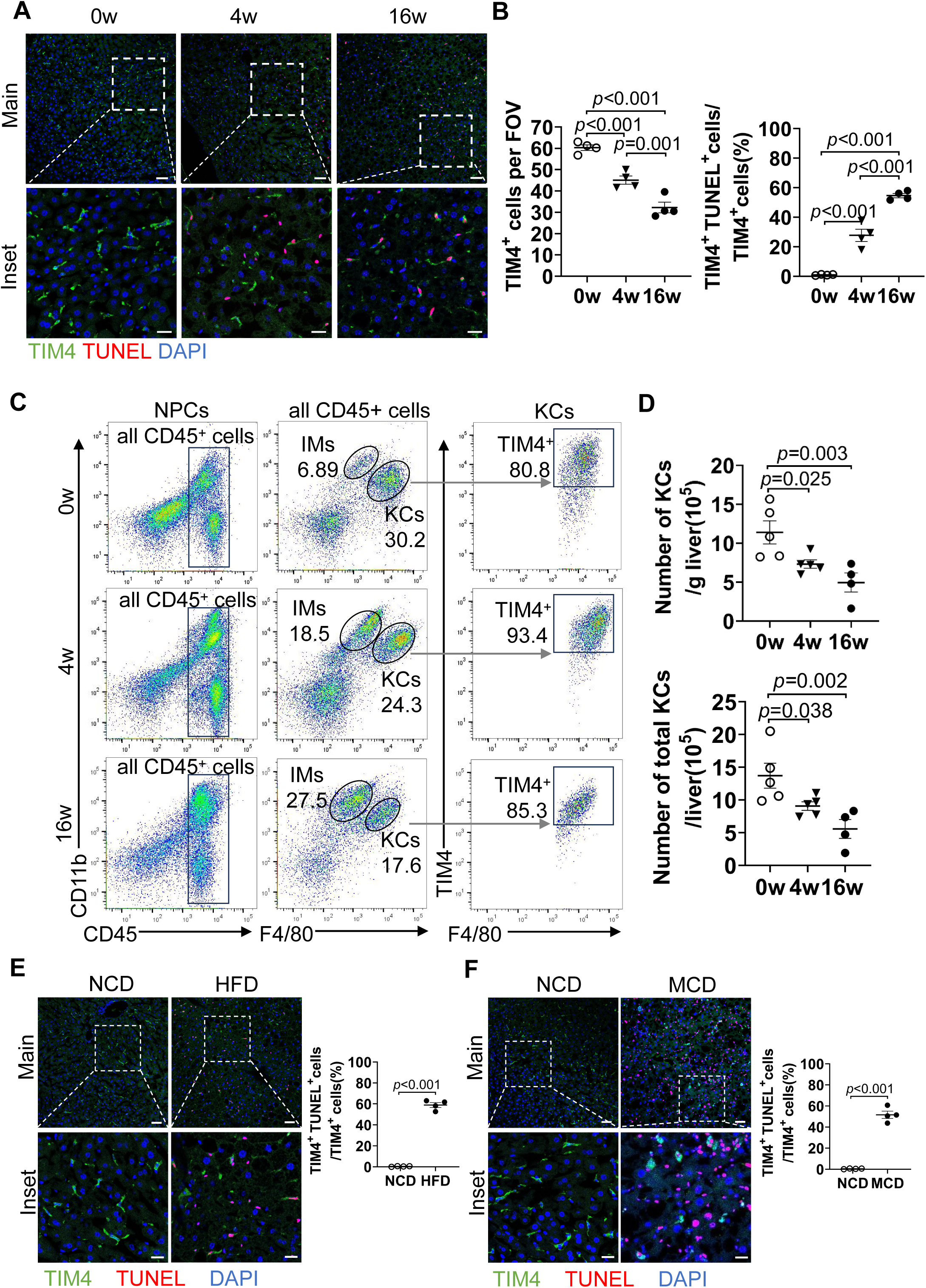
Kupffer cell death is a characteristic feature of MASLD progression **(A-D)** Male Wild-type C57BL/6J mice were fed a high-fat high-cholesterol diet (HFHC) for 0, 4, or 16 weeks. **(A)** KCs death was assessed by immunostaining of liver sections for TIM4(KCs marker, red), TUNEL (green), and DAPI (nuclei, blue). Scale bar: 50 µm (main panels) and 20 µm (Inset). **(B)** KCs death was quantified. n=4 mice/group. **(C)** Flow cytometry analysis of KCs (CD45^+^ F4/80^hi^ CD11b^low^ TIM4^+^) and IMs (CD45^+^ F4/80^low^CD11b^hi^ TIM4^-^) among isolated NPCs. **(D)** KCs counts were quantified. n=4-5 mice/group. **(E-F)** Male wild-type C57BL/6J mice were fed either: **(E)** Normal chow diet (NCD) or high-fat diet (HFD) for 20 weeks, or **(F)** NCD or methionine-choline-deficient diet (MCD) for 6 weeks. KCs death was assessed by immunostaining of liver sections for TIM4(green), TUNEL (red), and DAPI (nuclei, blue). Scale bar: 50 µm (main panels) and 20 µm (Inset). KCs death was quantified. n=4 mice/group. Representative images are shown in A, C, E, F. One-way ANOVA (B, D). Unpaired Student’s t-test (E, F). P value as indicated.

To exclude TUNEL false positivity associated with proliferation, we evaluated Ki67 expression in dying KCs (TUNEL^+^TIM4^+^). We observed that proliferating KCs (Ki67^+^TIM4^+^) were infrequent throughout HFHC feeding, and less than 15% of TUNEL^+^TIM4^+^ KCs co-expressed Ki67. This indicates that the majority of TUNEL^+^ KCs represent true cell death events rather than proliferation-related artifacts (Figure S2D). To examine the broader occurence of this phenomenon in MASLD, we further assessed KCs death in several other dietary mouse models, including mice fed a high-fat diet (HFD) for 20 weeks^24^ and a methionine/choline deficient diet (MCD) for 6 weeks^12^. Co-staining of TIM4 and TUNEL in liver sections revealed a notable increase in KCs death in both the HFD and MCD groups compared to the control group fed a normal chow diet (NCD) (Figure 1E, 1F). These findings collectively confirm that progressive KCs death is a pathological hallmark of MASLD observed across various dietary-induced models.

#### Kupffer cells exhibit early and zone-specific susceptibility to death during MASLD

### progression

To compare relative susceptibility of different hepatic cell populations to MASLD-induced death, we performed TUNEL co-staining with lineage markers: HNF4α (hepatocytes), Desmin (hepatic stellate cells; HSCs), and Iba1 (Hepatic macrophages, including both KCs and MoMFs) (Figure 2A–C). Hepatocyte death showed minimal change during the initial 4 weeks but increased modestly by 16 weeks (Figure 2A). In contrast, HSCs and hepatic macrophages mortality rose progressively throughout MASLD progression, with significant increases detectable as early as 4 weeks (Figure 2B, 2C). Notably, the finding that 60% of TIM4^+^ KCs are TUNEL^+^, compared to 30% of total Iba1^+^ cells, further supports that resident KCs undergo death more readily than the broader macrophage pool, which includes MoMFs.

**Figure 2.**
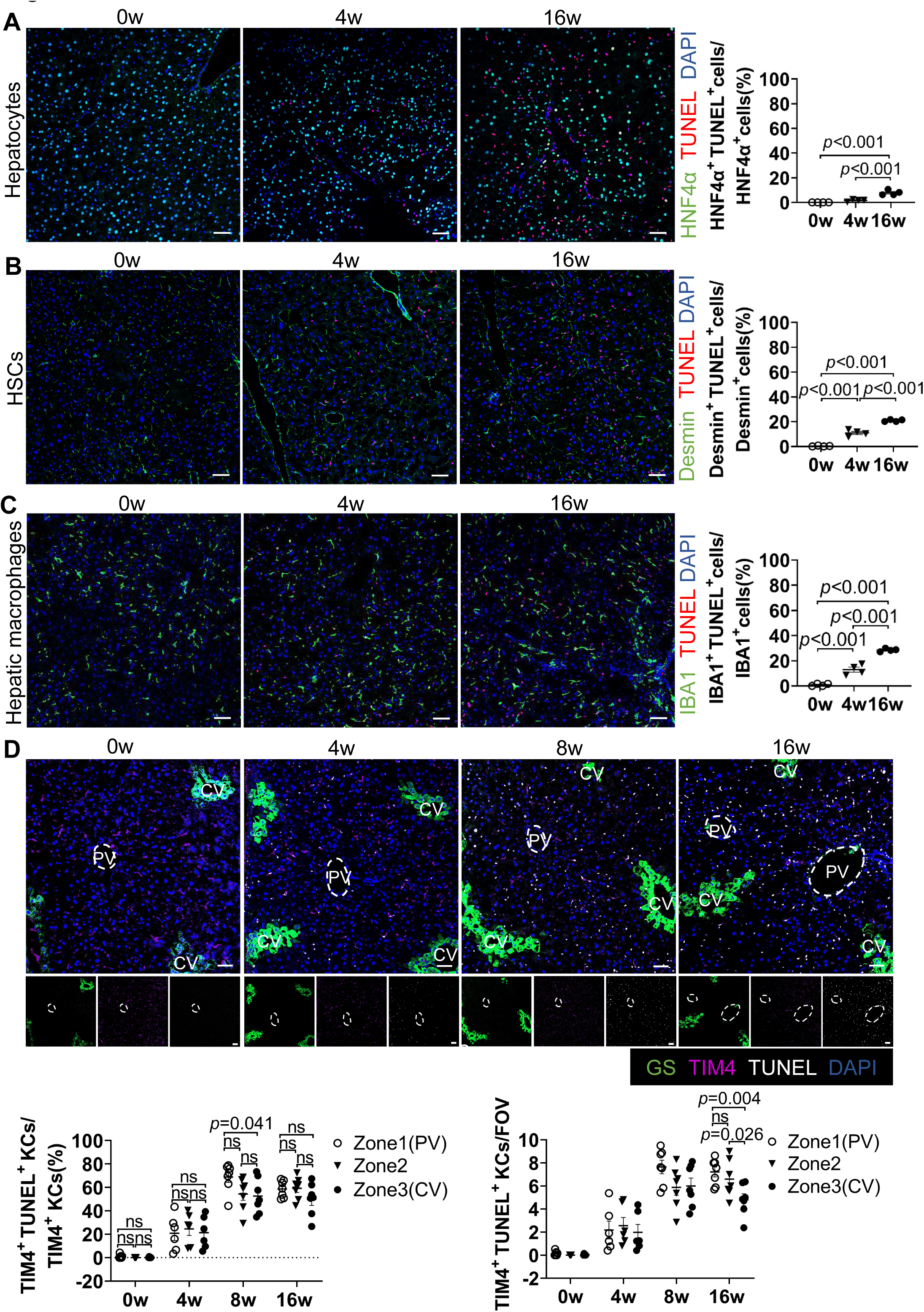
Kupffer cells exhibit early and zone-specific susceptibility to death during MASLD progression **(A-E)** Male Wild-type C57BL/6J mice were fed a HFHC diet for 0, 4, or 16 weeks. **(A-C)** Hepatic cell death was assessed by co-staining TUNEL with: **(A)** HNF4α (hepatocytes), **(B)** Desmin (hepatic stellate cells, HSCs), **(C)** Iba1 (hepatic macrophages), and DAPI (nuclei, blue). Scale bars: 50 µm (main panels). Hepatic cell death was quantified (n = 4 mice/group). **(D)** Zonal distribution of KCs death was evaluated by co-staining TIM4 (KCs), TUNEL, Glutamine Synthetase (GS, central vein marker) and DAPI (nuclei, blue). Scale bars: 50 µm. Zonal distribution of KCs death was quantified (n = 6-7 mice/group). FOV: field of view. PV: portal vein. CV: central vein. Representative images are shown in A-D. One-way ANOVA (A-D). P value as indicated.

We next determined whether KCs death displays spatial zonation within the liver lobule. Based on the lobular architecture—comprising periportal (PP, zone 1), midzonal (Mid, zone 2), and pericentral (PC, zone 3) regions^25^—we performed co-staining for TIM4, TUNEL, and glutamine synthetase (GS; a central vein marker) (Figure 2D). Spatial analysis revealed an early zonation bias in KCs death. At 8 weeks of HFHC feeding, the proportion of TIM4⁺TUNEL⁺cells among total TIM4⁺KCs was significantly higher in PV regions compared to CV regions (p = 0.041), indicating increased susceptibility of periportal KCs (Figure 2D). However, by 16 weeks, although absolute numbers of apoptotic KCs differed across zones, the proportional death rate became comparable (Figure 2D), suggesting that prolonged HFHC feeding leads to widespread KCs loss that attenuates early zonation bias.

#### Kupffer cells exhibit metabolic reprogramming with increased glycolysis during early MASLD

To define the glucose metabolic alterations that occur in KCs during MASLD development, we isolated KCs from wildtype mice at various time points of HFHC feeding (0, 4, 8, and 16 weeks). The purity of isolated KCs was confirmed by TIM4 immunofluorescence staining, with TIM4^+^ cells exceeding 90% (Figure S3A). We then conducted qRT-PCR to assess the mRNA expression levels of key enzymes involved in glycolysis (*Slc2a1, Hk3, Pfkfb3, Pkm*), the pentose phosphate pathway (PPP) (*G6pd, 6pdg*), glycogenolysis (*Pygl*), and glycogenesis (*Gys1, Ugp2*). Our data revealed that mRNA expression of rate-limiting enzymes for fast glucose metabolism, such as glycolysis and PPP, was significantly increased as early as 8 weeks after initiating the HFHC diet (Figure S3B). While glycolytic enzyme expression remained elevated, PPP enzyme expression began to decline by 16 weeks (Figure S3B). In contrast, enzymes linked to slower glucose metabolism, such as oxidative phosphorylation (*Idh1, Ogdh*), did not exhibit significant changes (Figure S3B). Furthermore, mRNA expression of glycogenesis and glycogenolysis rate-limiting enzymes started to decrease at 8 and 16 weeks, respectively, suggesting that glucose uptake becomes the primary source of KCs glucose metabolism during this period (Figure S3B). Additionally, we investigated mRNA expression of rate-limiting enzymes involved in β-oxidation (*Acadm, Hadh*) but found no significant differences (Figure S3B). These data suggest a time-dependent metabolic inflexibility, where impaired glycogen handling and mitochondrial inertia drive a sustained glycolytic dependence in KCs, which may exacerbate their vulnerability (particularly in periportal zones, where glycogen storage is predominant^26^).

To validate KCs-specific metabolic alterations in MASLD, we performed metabolomic analysis on primary KCs from wild-type mice fed an HFHC diet for 0, 4, or 8 weeks (Figure 3A). Given that KCs death peaked at 8 weeks in our model (Figure 2D), we focused on these early time points to capture initial metabolic shifts. Principal component analysis (PCA) of KCs metabolites revealed distinct, diet duration-dependent clustering (Figure 3B). Kyoto Encyclopedia of Genes and Genomes (KEGG) pathway enrichment analysis identified glucose metabolism pathways—including glycolysis and the pentose phosphate pathway (PPP)—as the most significantly upregulated during MASLD progression (Figure 3C, 3D). This glycolytic activation was further corroborated by time-dependent accumulation of key intermediates including glucose, Phosphatidylethylamine [PEA], Phosphoenolpyruvate [PEP], fructose-1,6-bisphosphate [FBP], and lactate [LA] in heatmap analysis (Figure 3E). Critically, we observed progressive increases in pro-apoptotic metabolites generated through these glucose metabolism pathways: redox disruptors (GSSG, FAD), mitochondrial toxins (methylmalonic acid), and apoptosis mediators (Hcy) - all exhibiting temporal coupling with glycolytic intermediates (Figure 3F). This demonstrates that KCs undergo rapid glycolytic reprogramming during early MASLD pathogenesis, which actively generates cytotoxic effectors coinciding with their peak vulnerability.

**Figure 3.**
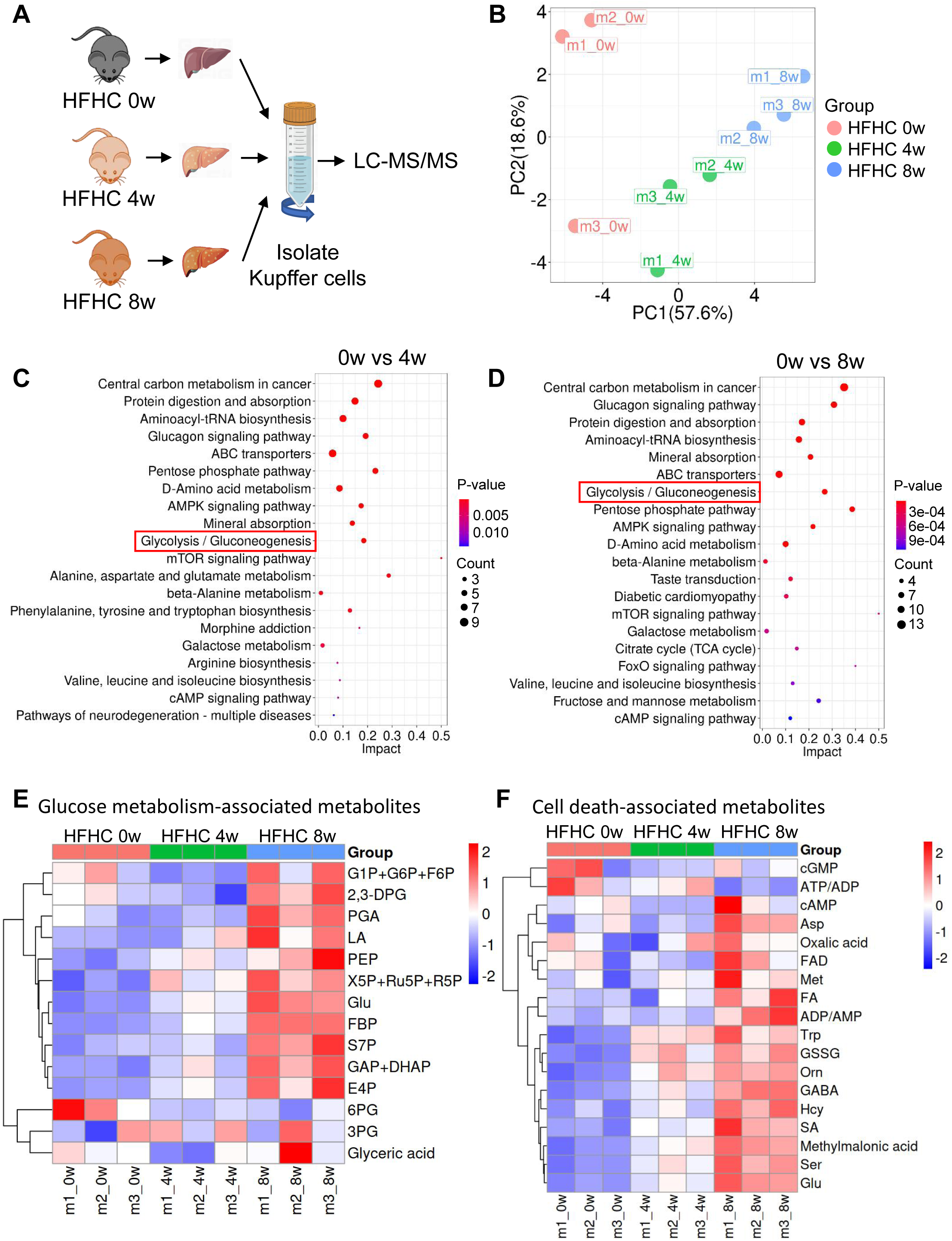
Kupffer cells exhibit metabolic reprogramming with increased glycolysis during early MASLD. **(A)** Experimental design for metabolomic analysis of KCs isolated from male wild-type mice fed a HFHC diet for 0, 4 or 8 weeks. n=3 mice/group. **(B)** Principal component analysis (PCA) of enriched metabolites in KCs across different dietary durations. **(C-D)** Kyoto Encyclopedia of Genes and Genomes (KEGG) pathway enrichment analysis of metabolic pathways upregulated in KCs at 4 weeks **(C)** or 8 weeks **(D)**. The glucose metabolism pathway is highlighted by red rectangles. **(E)** Heatmap depicting significantly altered metabolites involved in glucose metabolism pathways in KCs across different dietary durations. **(F)** Heatmap depicting significantly altered metabolites involved in cell death in KCs across different dietary durations.

#### Excessive glucose metabolic activity contributes to Kupffer cells death

To investigate the direct role of glucose metabolism in KCs death, isolated KCs were subjected to *in vitro* metabolic perturbations. First, KCs were treated with a combination of high glucose and palmitic acid (PA) to model MASLD pathology. Physiological glucose (5.5 mM) combined with PA (800 µM) increased KCs death by approximately 10%. Strikingly, glucose at a concentration mimicking HFHC feeding (10 mM) combined with PA (800 µM) increased KCs death by approximately 27%. This was evidenced by increased Cl-Casp3 staining (Figure 4A) and elevated Cl-Casp3 protein levels via western blot (Figure 4B).

**Figure 4.**
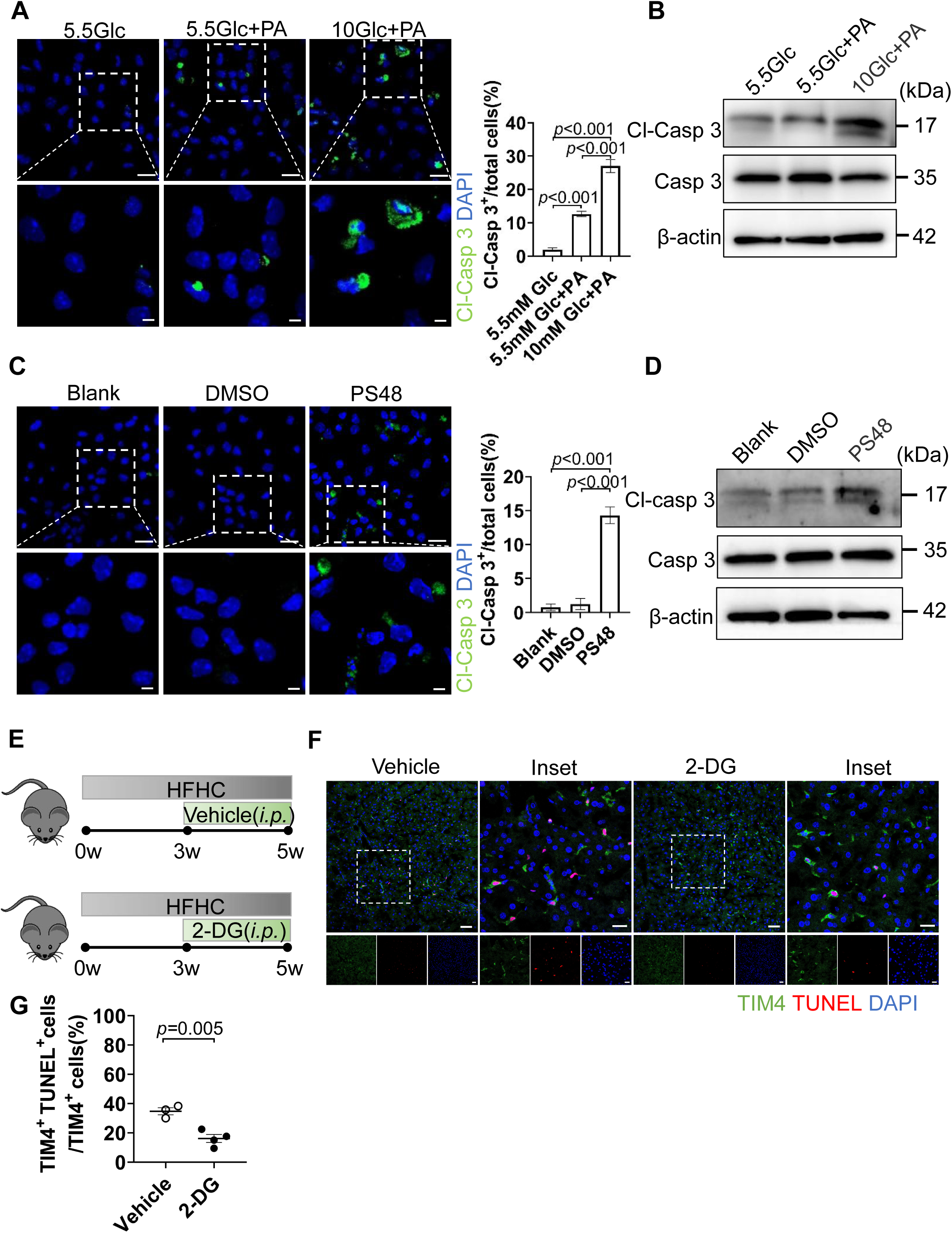
Excessive glucose metabolic activity contributes to Kupffer cell death. **(A-B)** Isolated KCs were treated for 24 h with: 5.5 mM glucose + isopropanol (control), 5.5 mM glucose + 800 µM palmitic acid (PA), 10 mM glucose + 800 µM PA. Cell viability was assessed by Cleaved caspase-3 (Cl-Casp3) staining (Cl-Casp3^+^ cells = dead). Scale bars: 20 µm (main panels), 5 µm (insets). Cl-Casp3 was detected by Western blot. **(C-D)** Isolated Kupffer cells were treated for 24 h with: Blank (no treatment), DMSO (vehicle control), 20 µM PS48 (PDK1 activator). Scale bars: 20 µm (main panels), 5 µm (insets). Cell death were analyzed as above. **(E)** Experimental design. Male WT mice were fed a high-fat, high-cholesterol (HFHC) diet for 5 weeks. From the 3rd week onward, mice received intraperitoneal injections of either vehicle or 2-DG (50 mg/kg) every other day (n = 3-4 mice/group). **(F-G)** Effects of glycolysis inhibition on KCs death after 5 weeks of HFHC feeding. (F) Representative images of liver sections co-stained with TUNEL and the KCs marker TIM4. (G) Quantification of TUNEL+ KCs. Data are presented as mean ± SEM. Statistical analysis was performed using one-way ANOVA (A, C) and an unpaired Student’s t-test (G); P values are indicated.

Second, we treated KCs with the Pyruvate Dehydrogenase Kinase (PDK1) activator PS48, which directly stimulates glycolysis. The results indicated a markedly increased KCs death compared to blank or DMSO vehicle controls (Figure 4C, 4D). Finally, Oligomycin (an ATP synthase inhibitor) was used to force glycolytic reliance by blocking mitochondrial ATP production. Oligomycin treatment similarly induced significant KCs death (Figure S4A, S4B), phenocopying the effects of direct glycolytic activation. KCs are difficult to culture and prone to death *in vitro*. To assess the health status of KCs in our metabolic perturbation assays, we also examined KCs viability using Calcein-AM staining, a fluorescent dye that labels metabolically active cells. Image analysis confirmed that the KCs were healthy and alive under all tested conditions (Figure S4C), indicating that our isolated KCs were suitable for *in vitro* experiments.

To determine whether enhanced glycolytic activity contributes to KCs death *in vivo* during MASLD, we pharmacologically inhibited glycolysis using 2-deoxy-D-glucose (2-DG). 2-DG functions as a glucose analog that enters cells but cannot be fully metabolized, thereby inhibiting glycolysis through blockade of hexokinase and phosphoglucose isomerase (Figure 4E). Wild-type C57BL/6J mice were fed a HFHC diet for five weeks to induce early-stage MASLD. To inhibit KCs glycolysis *in vivo*, mice received intraperitoneal injections of either vehicle or 2-DG (50 mg/kg) every other day beginning at week 3 of HFHC feeding. The relatively short duration of HFHC exposure (5 weeks) combined with low-dose 2-DG treatment was designed to minimize potential confounding effects of systemic glycolytic inhibition on hepatocytes. At the end of the five-week feeding period, KCs survival was assessed by co-immunostaining liver sections for the KCs marker TIM4 and TUNEL to detect apoptotic cells (Figure 4F). Quantitative analysis demonstrated that 2-DG-treated mice exhibited a significantly lower proportion of TUNEL^+^KCs compared with vehicle-treated controls (Figure 4G). These results indicate that pharmacological inhibition of glycolysis protects KCs from cell death during MASLD. Together with our *in vitro* data showing that glycolytic activation promotes KCs death.

#### Enhanced glycolytic flux in *Chil1^-/-^* macrophages

To investigate whether hyperactivated glycolysis specificly in KCs drives their death *in vivo*, we focused on Chitinase 3-like 1 (Chi3l1; gene *Chil1*)—a known inhibitor of glucose uptake^17^. Since Chi3l1 suppresses glucose uptake by KCs, KCs in *Chil1^-/-^* mice are expected to maintain a state of hyperactivated glycolysis. To confirm this, we first performed a glucose metabolic flux assay in KCs. However, due to the limited availability of primary KCs, we used bone marrow-derived macrophages (BMDMs) to dissect the Chi3l1-dependent metabolic mechanisms. Uniformly labeled [U-^13^C]glucose tracer analysis (Figure 5A) revealed genotype-specific metabolic reprogramming: PCA showed distinct clustering between *Chil1^-/-^* and wild-type (WT) BMDMs (Figure 5B), and heatmaps demonstrated pronounced accumulation of glycolytic intermediates (Figure 5C). Metabolic flux quantification confirmed significantly elevated U-^13^C enrichment in glycolytic metabolites—including glucose (Glc), fructose-6-phosphate (F6P), 3-phosphoglycerate (3PGA), 2-phosphoglycerate (2PGA), phosphoenolpyruvate (PEP), pyruvate (PA), and lactate (LA)—whereas glucose-6-phosphate (G6P), fructose-1,6-bisphosphate (FBP), and pentose phosphate pathway (PPP) intermediates (ribulose-5-phosphate [Ru5P], ribose-5-phosphate [R5P], sedoheptulose-7-phosphate [S7P]) remained unchanged. Dihydroxyacetone phosphate (DHAP) was the only PPP-linked metabolite showing increased enrichment (Figure 5D), indicating that *Chil1* deletion selectively hyperactivates glycolysis without engaging the PPP. Furthermore, extracellular acidification rates (ECAR) analysis also showed significantly elevated glycolytic capacity in *Chil1^-/-^* BMDMs (Figure 5E, 5F).

**Figure 5.**
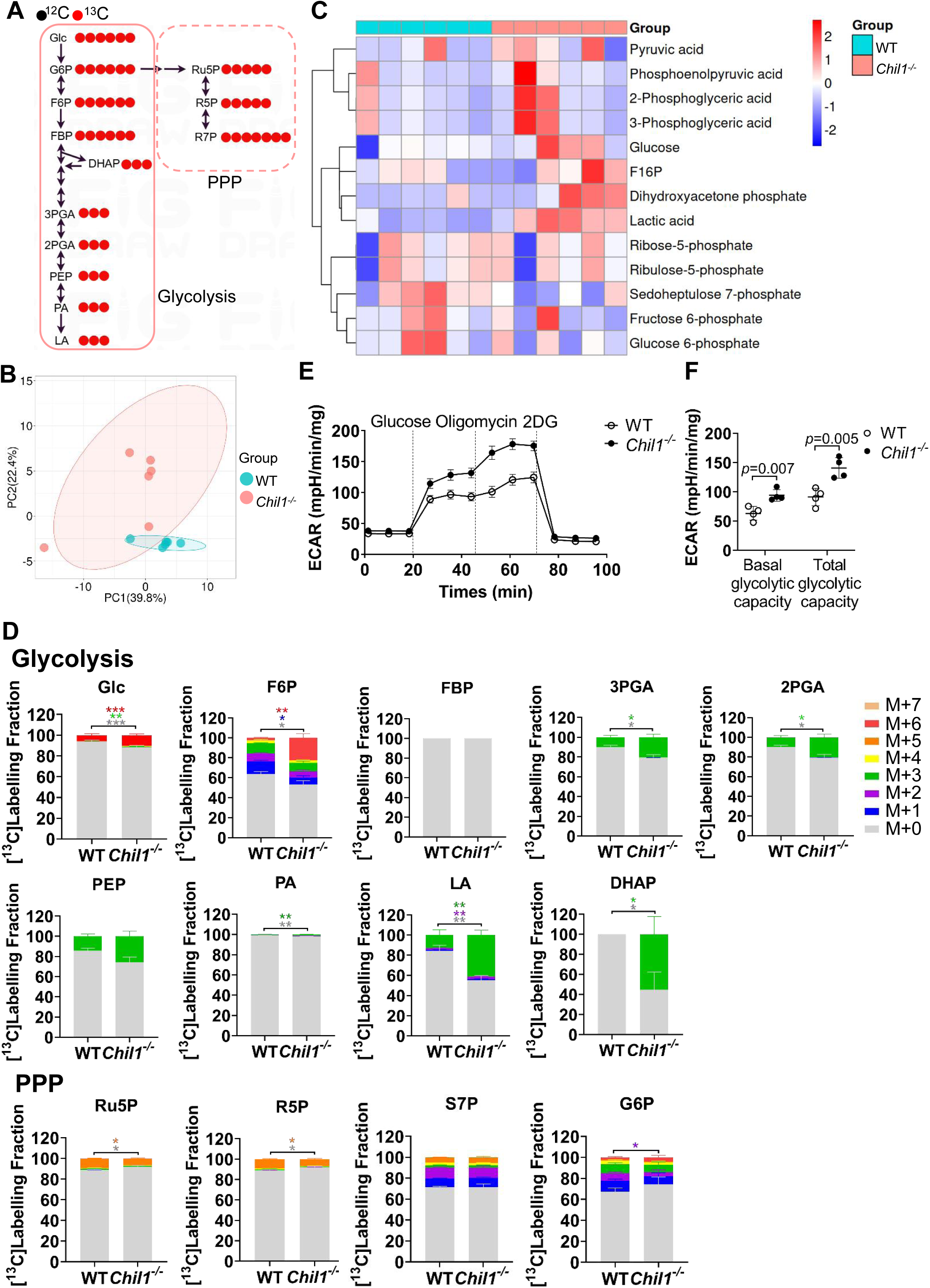
Enhanced glycolytic flux in *Chil1^-/-^* macrophages. **(A)** Schematic diagram depicting the fate of glucose-derived ribose carbons in WT mouse primary hepatocytes. **(B)** Principal component analysis (PCA) of metabolites in WT and *Chil1^-/-^* BMDMs cultured with [U-^13^C]glucose. **(C)** Heatmap depicting significantly altered Glycolysis and Pentose phosphate (PPP) metabolites in WT and *Chil1^-/-^* BMDMs. **(D)** Glucose metabolic flux analysis in WT and *Chil1^-/-^* BMDMs cultured with [U-^13^C]glucose showing mass isotopologue distributions of: Glycolytic intermediates (Glc, F6P, FBP, 3PGA, 2PGA, PEP, PA, LA, G6P). PPP intermediates (Ru5P, R5P, S7P, DHAP). Data represent n= 6 biological replicates/group. **(E-F)** Extracellular acidification rate (ECAR) analysis of WT or *Chil1^-/-^* BMDMs cells. BMDM were sequentially treated with Glucose, oligomycin and 2-DG as indicated during seahorse. Unpaired student t-test (D,F). P*<0.05, P**<0.01, P***<0.001.

Increased glycolysis is often associated with proinflammatory polarization of macrophages^15^. To assess the effect of Chil1 loss on KCs polarization, we further isolated KCs from WT and *Chil1^-/-^*mice fed a HFHC diet at 0, 8 and 16 weeks, and measured the mRNA expression of proinflammatory markers (*Nos2, Cxcl9, CIITA, CD86, Ccl3,* and *Ccl5*) and anti-inflammatory markers (*Chil3, Retnla, Arg1,* and *Mrc1*). In WT KCs, the mRNA levels of proinflammatory markers increased with MASLD progression: *Nos2, Cxcl9, CIITA,* and *Ccl3* rose significantly, while *CD86, Ccl5* showed a upward treand (Figure S5A). In contrast, *Chil1^-/-^* KCs exhibited higher baseline expression of proinflammatory genes, with *Nos2* and *Ccl5* significantly elevated and *Cxcl9* and *CD86* showing an increasing tendency compared to WT. As MASLD developed, proinflammatory gene expression increased further in *Chil1^-/-^*KCs. No significant differences in anti-inflammatory marker expression were observed between WT and *Chil1^-/-^* KCs at baseline. However, with MASLD progression, *Chil1^-/-^* KCs displayed a rising trend in the expression of anti-inflammatory markers, including *Chil3, Arg1,* and *Mrc1* (Figure S5A). Together, these data suggest that Chi3l1 deficiency drives KCs toward a partially pro-inflammatory phenotype.

Since Chi3l1 primarily functions in its secretory form as a macrophage glucose uptake inhibitor^17^, we also added recombinant Chi3l1(rChi3l1) supplementation group. rChi3l1 supplementation reversed the hyper-glycolytic flux, restoring it to levels comparable to those in WT macrophages (Figure S6A-S6D). This was accompanied by reduced mass isotopologue distributions of key glycolytic intermediates (Glc, F6P, FBP, 3PGA, 2PGA, PEP, PA, LA, G6P) (Figure S6B, S6C). In contrast, PPP intermediates (Ru5P, R5P, S7P, DHAP) were unaffected (Figure S6B, S6C), confirming the specificity of the effect on glycolysis. Functionally, rChi3l1 significantly lowered lactate dehydrogenase (LDH) activity in high glucose-treated BMDMs, further corroborating the attenuation of glycolytic output (Figure S6D). We acknowledge that detailed metabolic flux analysis was performed in BMDMs rather than primary KCs due to the technical challenge of obtaining sufficient viable cells for such assays. While ontogenically distinct from KCs, BMDMs are a widely accepted model for mechanistic metabolic dissection. Importantly, the hyperglycolytic phenotype observed in *Chil1^-/-^* BMDMs was consistent with our *in vivo* KCs data: Chi3l1 deficiency drove pro-inflammatory polarization. Thus, despite this limitation, the concordance between our *in vitro* mechanistic findings and *in vivo* phenotypic data provides compelling evidence that *Chil1^-/-^* mice serve as a mechanistic model to investigate glycolysis-driven KCs death during MASLD. Collectively, these data establish that Chi3l1 reduces glycolytic flux—without affecting PPP activity—in macrophages, thereby providing a mechanistic model to investigate glycolysis-driven KC death.

#### Enhanced glycolysis drives Kupffer cell death in MASLD

We therefore employed *Chil1^-/-^* mice as a mechanistic model to investigate glycolysis-driven KCs death. First, we isolated primary KCs from WT and *Chil1^-/-^* mice and treated them with PA or Iso, or left untreated (Blank). *Chil1^-/-^* KCs exhibited significantly increased susceptibility to PA-induced cell death compared to WT controls *in vitro,* as indicated by elevated Cl-Casp3 immunostaining (Figure 6A, 6B). This result was further confirmed by increased LDH release from *Chil1^-/-^*KCs (Figure 6C). To delineate the cellular source of Chi3l1 in MASLD livers, we performed immunostaining on serially sectioned liver tissues from mice fed an HFHC diet for 16 weeks. Consecutive sections were independently probed for Chi3l1 or lineage-specific markers (HNF4α, Desmin, Iba1), enabling cellular localization through morphological alignment across sequential slices. This revealed predominant Chi3l1 expression in hepatic macrophages (Figure S7A). Moreover, Chi3l1 expression was significantly elevated in KCs (F4/80^+^TIM4^+^) of HFHC-fed mice, compared to NCD controls (Figure S7B), suggesting Chi3l1 is primarily expressed in KCs and may function as a KCs-autonomous regulator.

**Figure 6.**
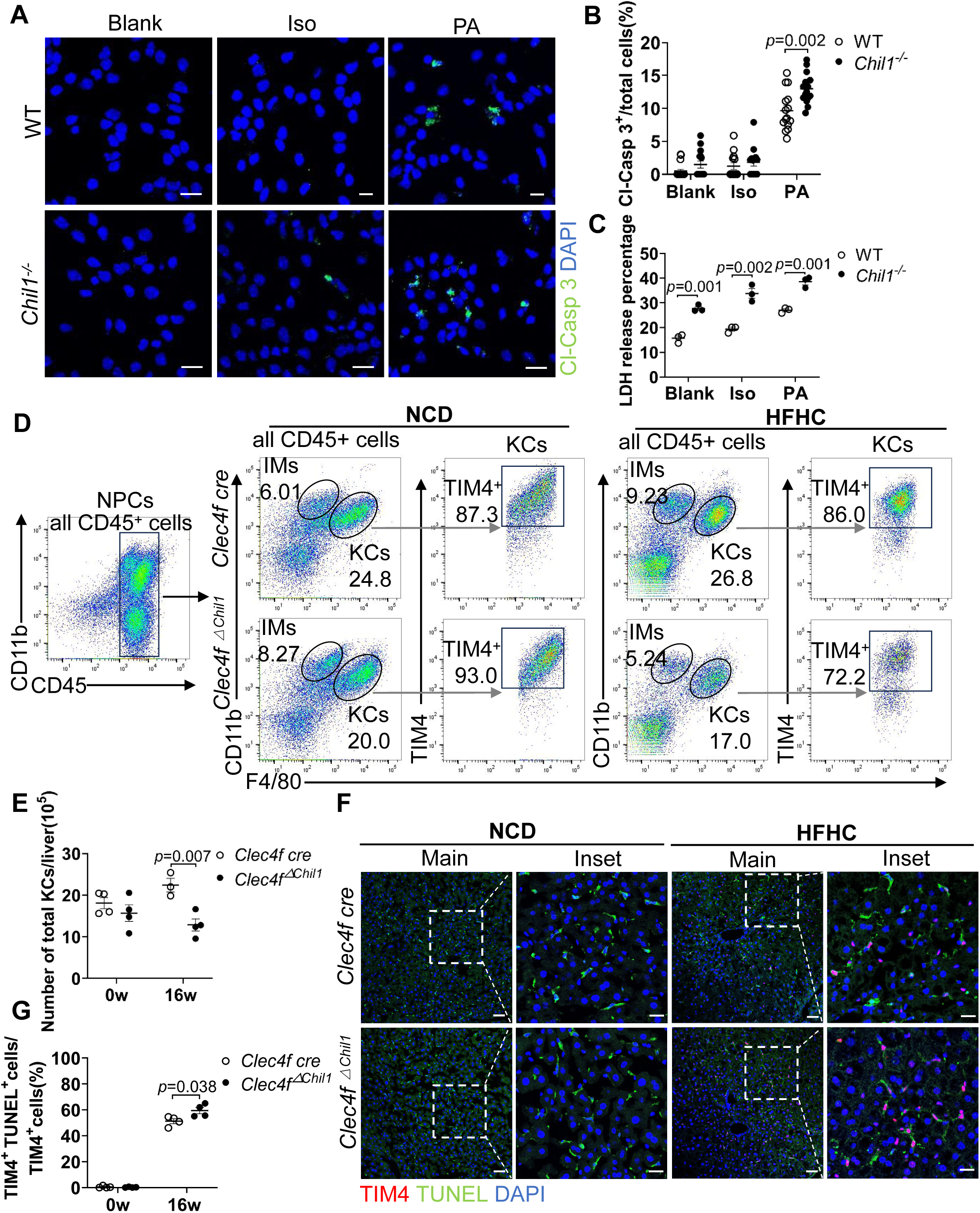
Enhanced glycolysis accelerated Kupffer cell death during MASLD. **(A)** Cleaved caspase-3 (Cl-Casp3) staining to detect WT and *Chil1^-/-^* Kupffer cell death. Cells were under treatment without (Blank) or with either Isopropyl alcohol (Iso) or Palmitic Acid (PA) for 24 h. Scale bar: 20μm. **(B)** Cl-Casp3+ cells were quantified. **(C)** LDH release measurement in culture medium of KCs isolated from male WT and *Chil1^-/-^*mice was measured after treatment for 24 h with: Blank (no treatment), ISO (vehicle control), 800 µM PA. **(D)** Flow cytometry analysis of KCs (CD45^+^ F4/80^hi^ CD11b^low^ TIM4^+^) and MoMFs (CD45^+^ F4/80^low^ CD11b^hi^ TIM4^-^) among NPCs in *Clec4f-cre* and *Clec4f^ΔChil1^* mice fed HFHC diet for 0 or 16 weeks. **(E)** KCs counts were quantified. n= 4 mice/group. **(F)** Kupffer cell death was assessed by immunostaining of TIM4 (KCs marker, green), TUNEL (red), and DAPI (nuclei, blue) in liver sections from *Clec4f-cre* and *Clec4f^ΔChil1^* mice fed HFHC diet for 0 or 16 weeks. Scale bar: 50μm (main panels) and 20μm (Inset). **(G)** KCs death was quantified. n=4 mice/group. Representative images shown (A, D, F). Unpaired student t-test (B,C,E,G). P value as indicated.

**Figure 7.**
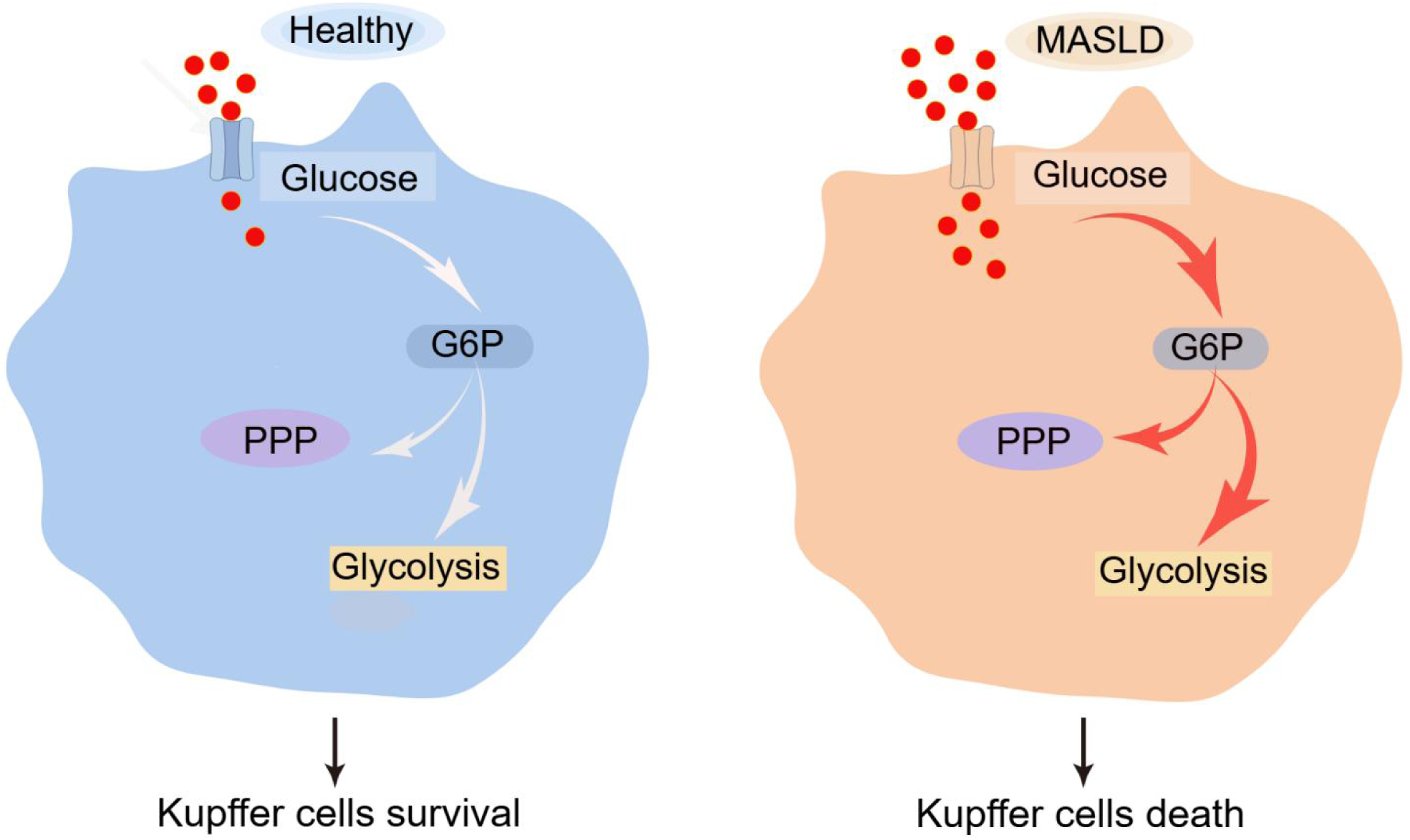
Excessive glycolysis enhancement promotes Kupffer cell death in MASLD. (Left) Under physiological conditions, Kupffer cells (KCs) maintain basal glucose metabolism supporting cellular homeostasis and survival. **(Right)** During MASLD progression, KCs undergo excessive glycolysis enhancement, which accelerates KCs death.

To determine cell-autonomous effects, we generated KCs-specific Chil1 knockout mice (*Clec4f^ΔChil1^* mice) and analyzed their response to HFHC feeding. Flow cytometry of NPCs revealed a significant lower KCs population (CD45^+^ F4/80^hi^ CD11b^low^ TIM4^+^) in *Clec4f^ΔChil1^*mice compared to *Clec4f cre* controls after 16 weeks of HFHC diet (Figure 6D, 6E). During this analysis, we observed no reduction in KCs in *Clec4f cre* control mice, raising the possibility that Cre insertion itself might influence KCs mainteinence. To test this, we performed co-staining for TIM4 and Ki67, which revealed significantly higher numbers of proliferating KCs in *Clec4f cre* mice compared with C57BL/6J mice under NCD. Notably, following HFHC feeding, *Clec4f cre* mice exhibited an even greater increase in KCs proliferation (Figures S8A and S8B), suggesting that cre insertion enhanced KCs self-renewal which contributes to maintenance of the KCs pool in these mice under MASLD. Despite this proliferative baseline, *Clec4f^ΔChil1^* mice still displayed a reduced KCs number (Figure 6D, 6E) and increased KCs death. This was corroborated by TIM4/TUNEL co-staining, which showed elevated KCs death in *Clec4f^ΔChil1^*livers (Figure 6F, 6G). Collectively, these findings demonstrate that genetic ablation of *Chil1* amplifies glycolytic flux and sensitizes KCs to lipotoxic stress *in vitro*, and drives KCs depletion through accelerated cell death *in vivo*. Although Chi3l1 is not exclusively expressed by KCs, these data establish it as a critical regulator of KCs survival in MASLD, likely acting in a cell-autonomous manner within the KCs population.

## Discussion

This study identifies KCs death as an early and selective pathological feature of MASLD across multiple dietary models. Compared with other hepatic cell populations, KCs exhibit markedly increased susceptibility to apoptosis during early disease. Through metabolomic and functional analyses, we demonstrate that KCs undergo progressive metabolic reprogramming characterized by sustained activation of glycolysis. Importantly, our data support a causal role for excessive glycolytic flux in driving KCs death. Pharmacologic glycolytic stimulation (high glucose or PDK1 activation with PS48), enforced glycolytic dependence (oligomycin), and KC-specific Chi3l1 deletion each accelerated KC apoptosis, whereas glycolytic inhibition (2-DG) was protective. Together, these findings establish hyperactivation of glycolysis as a central mechanism underlying KCS vulnerability and depletion in MASLD (Figure 7).

KCs loss has been reported to varying degrees across MASLD models, reflecting differences in diet composition, disease duration, macrophage heterogeneity, and marker selection. For example, in a 6-week MCD diet model, approximately 10% of CLEC4F⁺ KCs were TUNEL⁺, indicating apoptotic death^12^. In contrast, another 6-week MCD study reported a marked reduction in TIM4⁺ KCs, from 66% to 26%^27^. More moderate KCs loss has been observed in HFD models, with a ∼20% reduction of TIM4⁺ KCs after 16 weeks^14^, whereas prolonged western diet feeding resulted in a >50% decrease in TIM4⁺ KCs at 36 weeks^14^. Together, these studies underscore the model-dependent nature of KCs loss and highlight the importance of experimental context and marker selection when assessing KCs dynamics in MASLD. Importantly, our spatial analyses further reveal preferential periportal KCs death during early MASLD. Given the metabolic zonation of the liver and higher periportal glucose flux, this regional susceptibility supports a mechanistic link between glucose metabolism and KCs survival. These findings suggest that metabolic microenvironment, in addition to inflammatory signaling, shapes KCs fate during MASLD progression.

While glycolytic reprogramming is classically associated with macrophage activation^15^, our data reveal a critical distinction in tissue-resident KCs. In MASLD, sustained and excessive glycolytic activation does not simply maintain inflammatory function but instead drives apoptotic loss. Thus, glycolysis in this context becomes maladaptive, serving as a determinant of cell fate rather than merely an activation program. This shifts the conceptual framework from inflammation-centered KCs dysfunction to metabolism-driven KCs attrition. Our findings also clarify the context-dependent role of Chi3l1 in liver disease. Prior studies in advanced MASH and fibrotic models have described pro-fibrotic functions of Chi3l1, including activation of hepatic stellate cells and modulation of macrophage survival pathways^28,29,30^. In contrast, using an HFHC model that recapitulates steatohepatitis without fibrosis, we identify a distinct early-stage role for Chi3l1 in maintaining KCs metabolic homeostasis. KC-specific deletion of Chi3l1, but not broader myeloid deletion, accelerated KCs loss and worsened steatosis^17^. These results establish Chi3l1 as a KCs-autonomous regulator that restrains glycolytic overload. The apparent duality of Chi3l1 function likely reflects disease stage and cellular source: protective in early MASLD through preservation of resident KCs, but pro-fibrotic in advanced disease when derived from monocyte-derived macrophages and acting on stellate cells.

Our prior work demonstrated that KCs exhibit relative metabolic inflexibility compared with monocyte-derived macrophages, relying heavily on glucose utilization^17^. The current study extends this concept by showing that loss of Chi3l1 leads to excessive glycolytic flux that exceeds KCs metabolic tolerance as supported by ¹³C-glucose tracing, enhanced lipotoxic sensitivity *in vitro*, and accelerated cell death *in vivo*. Thus, Chi3l1 functions as a metabolic checkpoint that fine-tunes glucose uptake and prevents lethal glycolytic overload. This selective dependence explains why Chi3l1 deficiency disproportionately affects KCs while sparing other macrophage populations.

Several limitations merit consideration. First, although multiple murine dietary models were used, validation in human MASLD—particularly spatial assessment of KC death and metabolic signatures—is necessary. Second, while we identify glycolytic hyperactivation as an upstream driver, the downstream apoptotic mechanisms remain incompletely defined. Potential contributors include oxidative stress, redox imbalance, lactate accumulation, or mitochondrial insufficiency under high glycolytic flux. Third, the interplay between glucose metabolism and other metabolic pathways, including fatty acid oxidation, warrants further investigation.

Despite these limitations, our findings have translational implications. Therapeutic strategies that preserve KCs metabolic balance—rather than broadly suppressing inflammation—may represent a novel approach for MASLD. Modulation of Chi3l1 signaling or selective targeting of key glycolytic regulators within KCs could help maintain the resident macrophage pool, preserve hepatic immune homeostasis, and potentially delay disease progression.In summary, we identify excessive glycolytic reprogramming as a fundamental mechanism driving selective KC apoptosis in early MASLD, with periportal predominance. Chi3l1 functions as an endogenous metabolic safeguard that restrains this lethal shift. Targeting KCs-specific metabolic vulnerability may offer a new strategy to preserve hepatic immune integrity and modify MASLD progression.

## Abbreviations

MASLD, metabolic dysfunction-associated fatty liver disease; MASH, metabolic dysfunction-associated steatohepatitis; KCs, Kupffer cells; Chi3l1, Chitinase 3 like 1; ALT, Alanine aminotransferase; AST, aspartate aminotransferase; TC, cholesterol; TG, triglyceride; NPCs, nonparenchymal cells; HFHC, high fat high cholesterol diet; NCD, normal chow diet; Clec4f, C-type lectin domain family 4; TIM4,T cell immunoglobulin mucin protein 4; MoMFs, monocyte-derived macrophages; HFD, high-fat diet; MCD, methionine/choline deficient diet; WD, western diet; PPP, pentose phosphate pathway; BMDM, bone marrow derived macrophages; DMSO, dimethyl sulfoxide; MAFL, non-alcoholic fatty liver; rChi3l1, recombinant murine Chi3l1; PA, palmic acid; Iso, Isopropyl alcohol; MoKCs, monocytes-derived Kupffer cells; ALD, alcohol-induced liver disease; AILI, acetaminophen-induced liver injury; EmKCs, embryo-derived Kupffer cells; DT, diphtheria toxin; WT, wild-type; TUNEL, TdT-mediated dUTP Nick-End Labeling; 2-DG, 2-deoxy-D-glucose.

## Acknowledgements

We thank Dr. Bin Qi (Yunnan University) for suggestions and discussion. We thank Dr. Guangxun Meng (Hainan Academy of Medical Sciences) for providing us with L929 cells. We thank Dr. Cynthia Ju (UTHealth,Texas) for advice in manuscript submission.

## Author Contributions

JH performed the experiments, analyzed the data, and wrote the manuscript. RL conducted immunofluorescence staining for zonal distribution analysis. CX and XEZ participated in sample collection. KQW assisted with metabolomics analysis. ZS conceived and designed the study, supervised the project, and wrote the manuscript.

---

**Figure S1.**
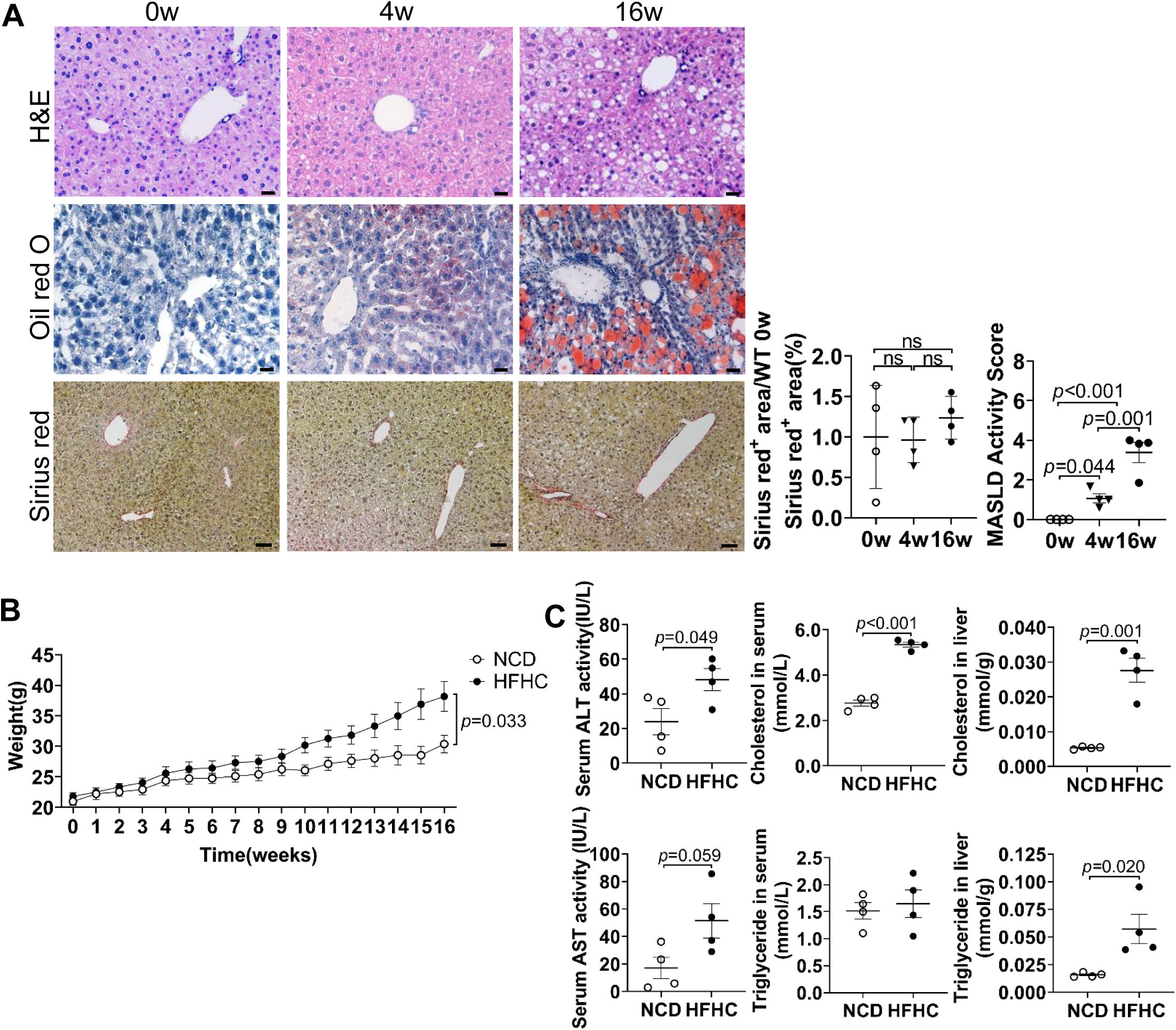
The generation of HFHC-induced MASLD mouse model. Male wildtype C57B/6J mice were fed with HFHC diet for 0, 4 or 16 weeks. **(A)** H&E (top), Oil red O (middle) and Sirius red staining (bottom) were performed to detect MASLD progression in liver sections at 0, 4 or 16 weeks after HFHC diet, respectively. Scale bar: 20μm. Liver fibrosis was quantified. MASLD Activity Score is diagnosed. **(B)** Body weight was recorded during HFHC diet feeding. **(C)** Serum ALT, AST, cholesterol, triglyceride or liver cholesterol, triglyceride is measured at 16 weeks after HFHC diet. n=4 mice/group. Unpaired Student’s t-test (B, C); one-way ANOVA (A). P value as indicated.ns: not significant.

**Figure S2.**
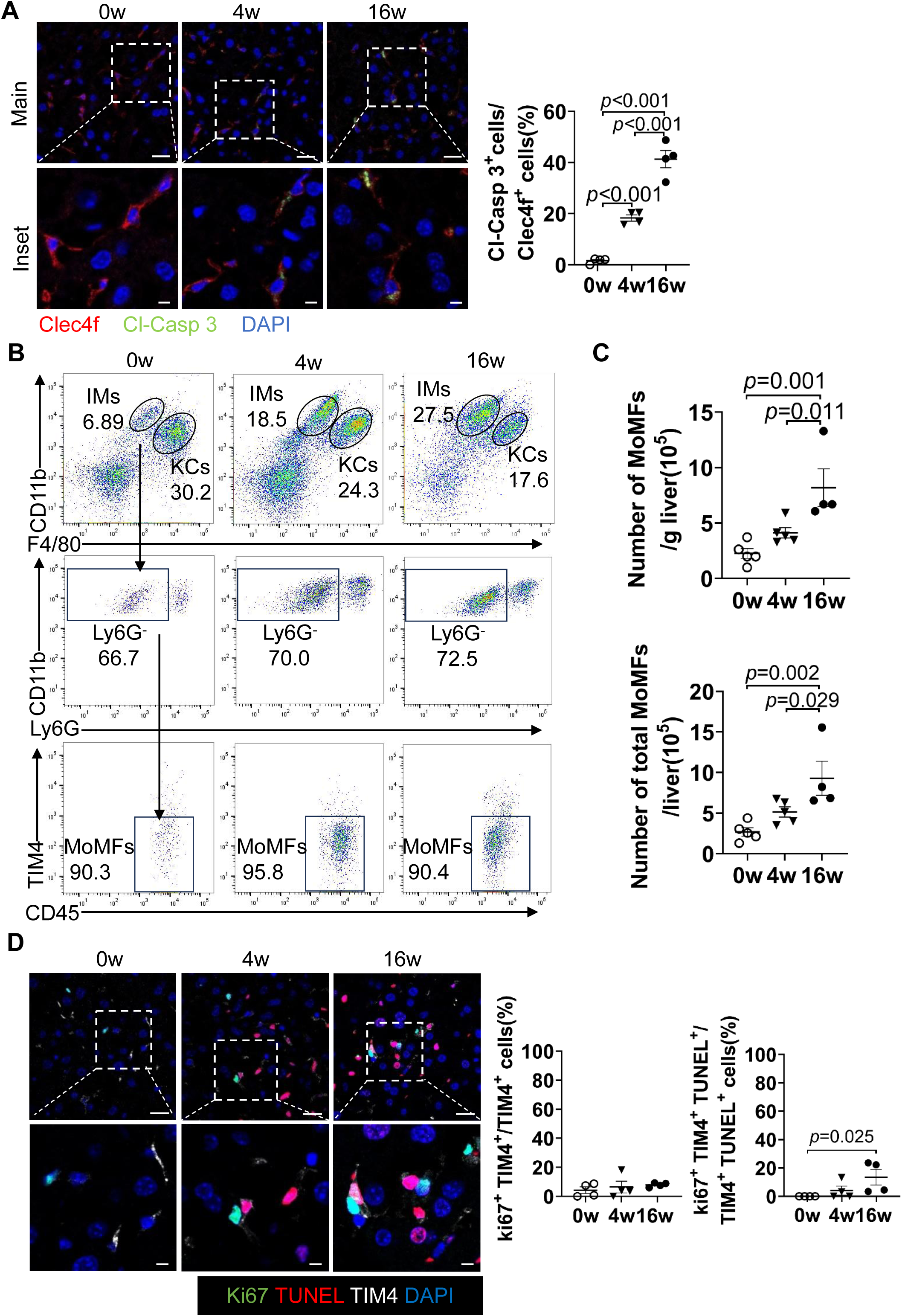
Examination of KCs death and MoMFs recruitment in HFHC mice. Male wildtype C57B/6J mice were fed with HFHC diet for 0, 4 or 16 weeks. **(A)** KCs death was examined by immunofluorescence staining of Clec4f and cleavaged caspase 3 (Cl-Casp3) and DAPI (Nuclei) in livers of mice. KCs death was quantified. n=4 mice/group. **(B)** Flow cytometry analysis of MoMFs (CD45⁺ Ly6G⁻ CD11b⁺ F4/80^low^TIM4^low/-^) among NPCs of WT mice fed HFHC deit for 0, 4, or 16 weeks. **(C)** MoMFs counts were quantified. n=4-5 mice/group. **(D)** Proliferation-associated TUNEL^+^ KCs was examined by co-staining of Ki67, TIM4, TUNEL and DAPI (Nuclei). Ki67^+^KCs among TUNEL^+^ KCs was quantified. Representative images are shown in A, D. One-way ANOVA (A, C, D). P value as indicated. Scale bar: 20μm (main panels), 5μm(inset).

**Figure S3.**
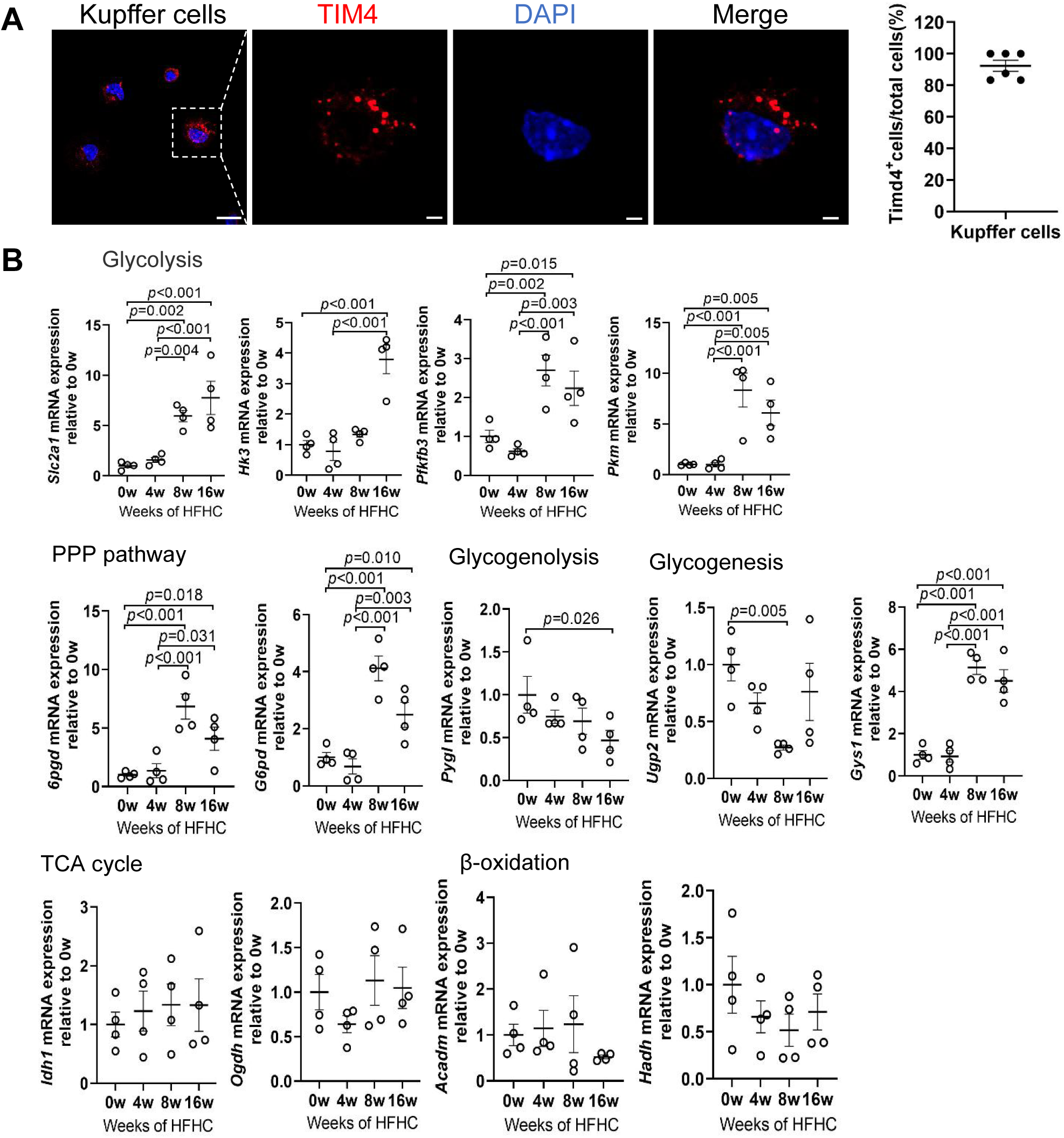
Dynamic changes in mRNA expression levels of rate-limiting enzyme genes involved in glucose metabolism. **(A)** Purity of isolated KCs was examined by immunofluorescence staining of TIM4 and DAPI (nuclei). Scale bar:20μm (main panels), 5μm(inset). Purity was quantified (n=6 independent experiments). **(B)** qRT-PCR analysis of mRNA expression levels of key rate-limiting enzymes in glycolysis, pentose phosphate pathway (PPP), glycogenolysis, glycogenesis, TCA cycle, and β-oxidation in WT KCs from mice for indicated dietary durations (n = 4 mice/group).One-way ANOVA (B), P values as indicated.

**Figure S4.**
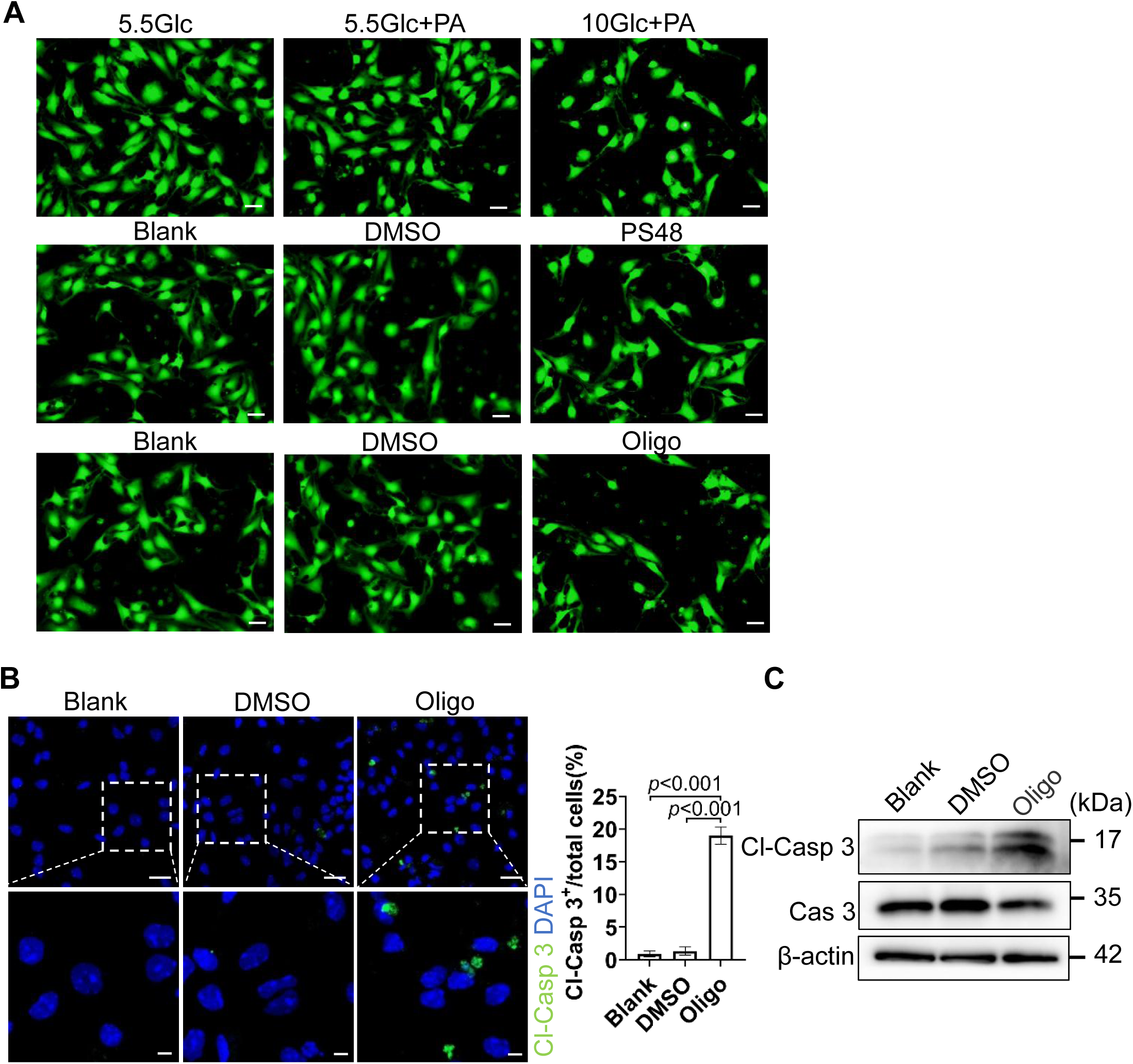
Excessive glucose metabolic activity contributes to Kupffer cell death. **(A)** Calcein-AM was used to assess primary KCs cells viability treated as in main Figure 4. Scale bar: 20μm. Representative images are shown. **(B-C)** Isolated Kupffer cells were treated for 24 h with: Blank (no treatment), DMSO (vehicle control), 20 µM oligomycin (Oligo, ATP synthase inhibitor). Scale bars: 20 µm (main panels), 5 µm (insets). Cell death were analyzed as above. One-way ANOVA (B). P value as indicated.

**Figure S5.**
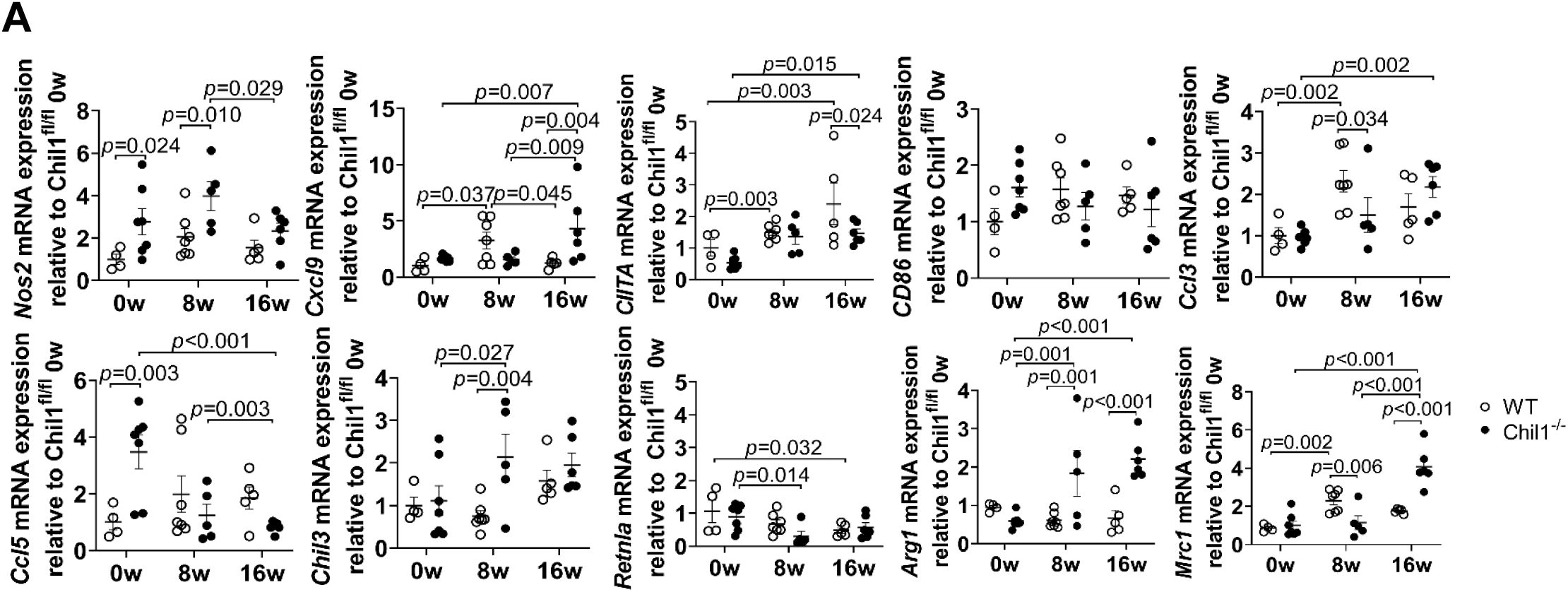
Comparison of KCs polarization between WT and *Chil1^-/-^* mice during MASLD progression . **(A)** qPCR analysis of Key genes involved in macrophage polarization pathway in liver tissues of WT and *Chil1^-/-^* mice fed with HFHC at indicated week. n=4-7 mice/group. Representative images are shown in A. One-way ANOVA (A). P value as indicated.

**Figure S6.**
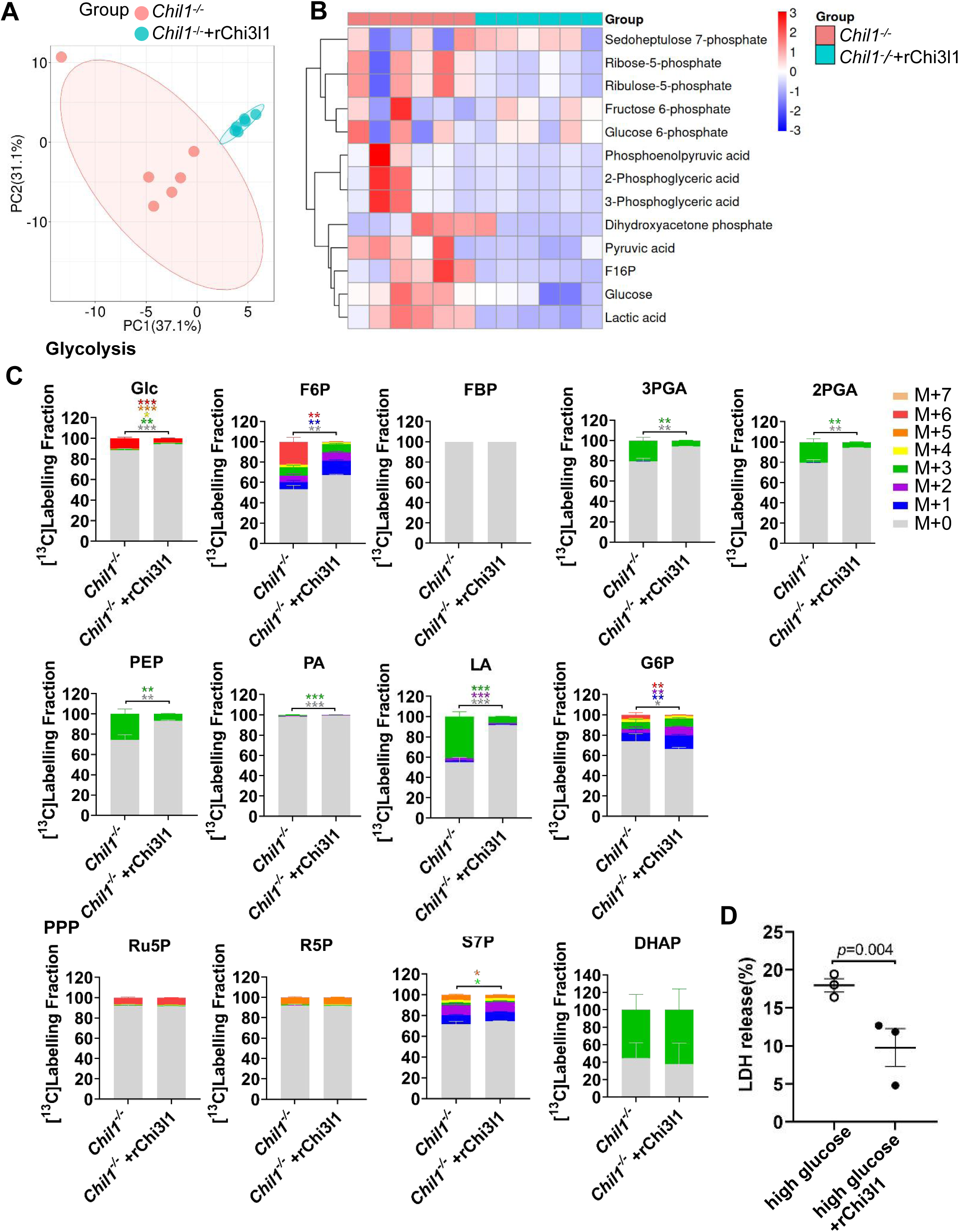
Recombinant Chi3l1(rChi3l1) inhibits glucose utilization in *Chil1^-/-^* BMDMs. **(A)** Comparative principal component analysis (PCA) of metabolites in *Chil1^-/-^*and *Chil1^-/-^* supplemented with rChi3l1 BMDMs cultured with uniformly labeled [U-^13^C]glucose. **(B)** Heatmap depicting significantly altered Glycolysis and Pentose phosphate (PPP) metabolites in *Chil1^-/-^* and *Chil1^-/-^* supplemented with rChi3l1 BMDMs . **(C)** Glucose metabolic flux analysis in *Chil1^-/-^* and *Chil1^-/-^* supplemented with rChi3l1 BMDMs showing mass isotopologue distributions of: glycolysis pathway intermediates (Glc, F6P, FBP, 3PGA, 2PGA, PEP, PA, LA, G6P). Pentose phosphate pathway intermediates (Ru5P, R5P, S7P, DHAP). Data represent n= 6 biological replicates. **(D)** Lactate dehydrogenase (LDH) activity in culture medium of *Chil1^-/-^*BMDMs treated for 24 h with: 10 mM glucose (high glucose), 10 mM glucose + 100 ng/mL rChi3l1. Unpaired Student’s t-test (D). P value as indicated.

**Figure S7.**
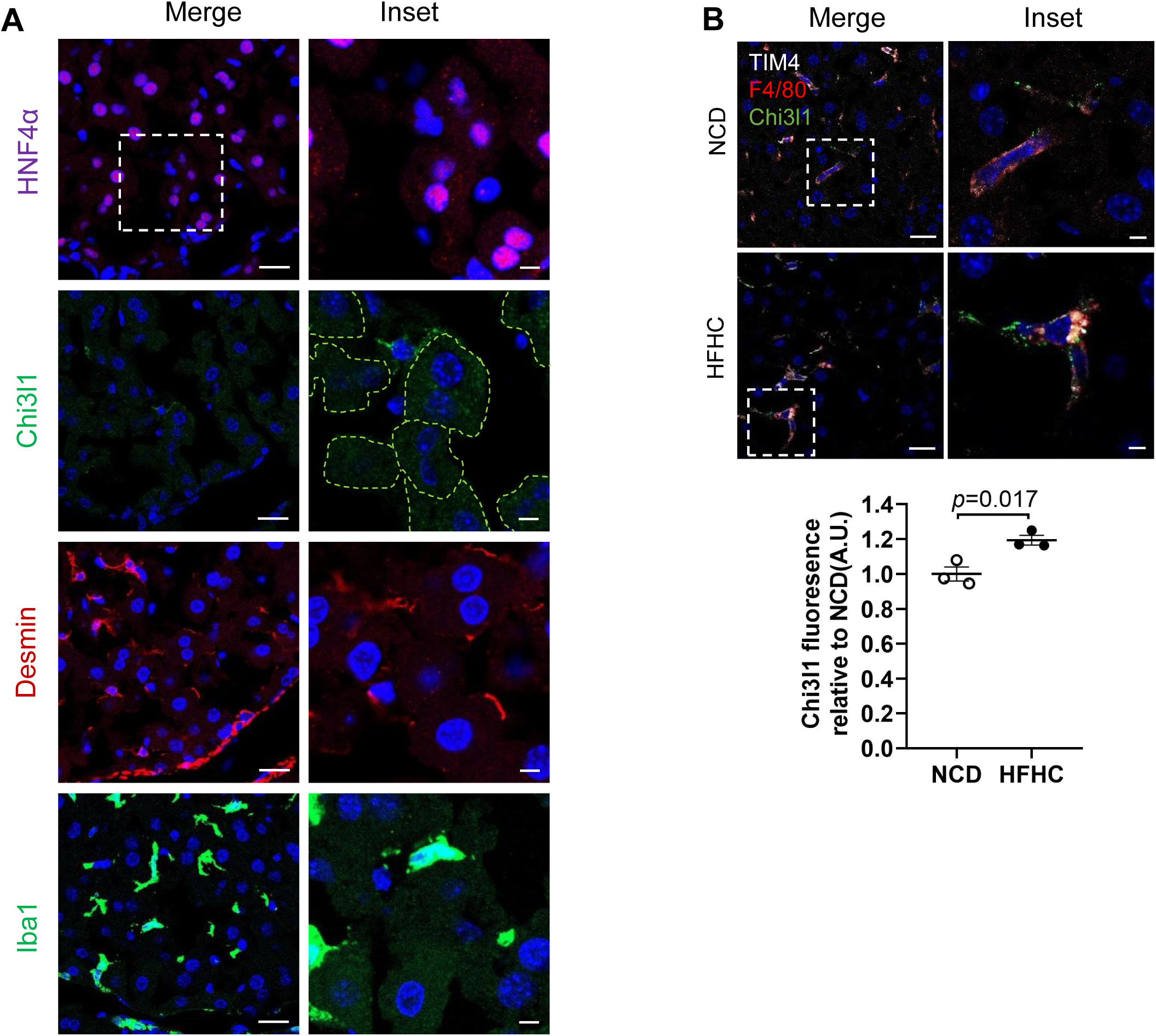
Chi3l1 is majorly expressed in hepatic macrophages and its expression is upreguated during MASLD. **(A)** Cellular source of Chi3l1 was assessed by immunohistochemical analysis of consecutive liver sections from mice fed an HFHC diet for 16 weeks. Serial sections were independently stained for: Chi3l1, Lineage markers: HNF4α (hepatocytes), Desmin (hepatic stellate cells; HSCs), or Iba1 (Hepatic macrophages). Cellular localization was determined by aligning morphological profiles across sequential sections. Scale bars: 20 μm (main panels), 5 μm (insets). Hepatocytes are outlined in Chi3l1+ images in green dashed lines. **(B)** Chi3l1 expression in KCs were assessed by co-staining with F4/80 (red) and Timd4 (white) in livers of mice fed either normal chow diet (NCD, 0 weeks) or HFHC diet (16 weeks). Scale bar:20μm (main panels), 5μm (inset). The intensity of Chi3l1 expression in KCs was quantified. Unpaired Student’s t-test (B). P value as indicated.

**Figure S8.**
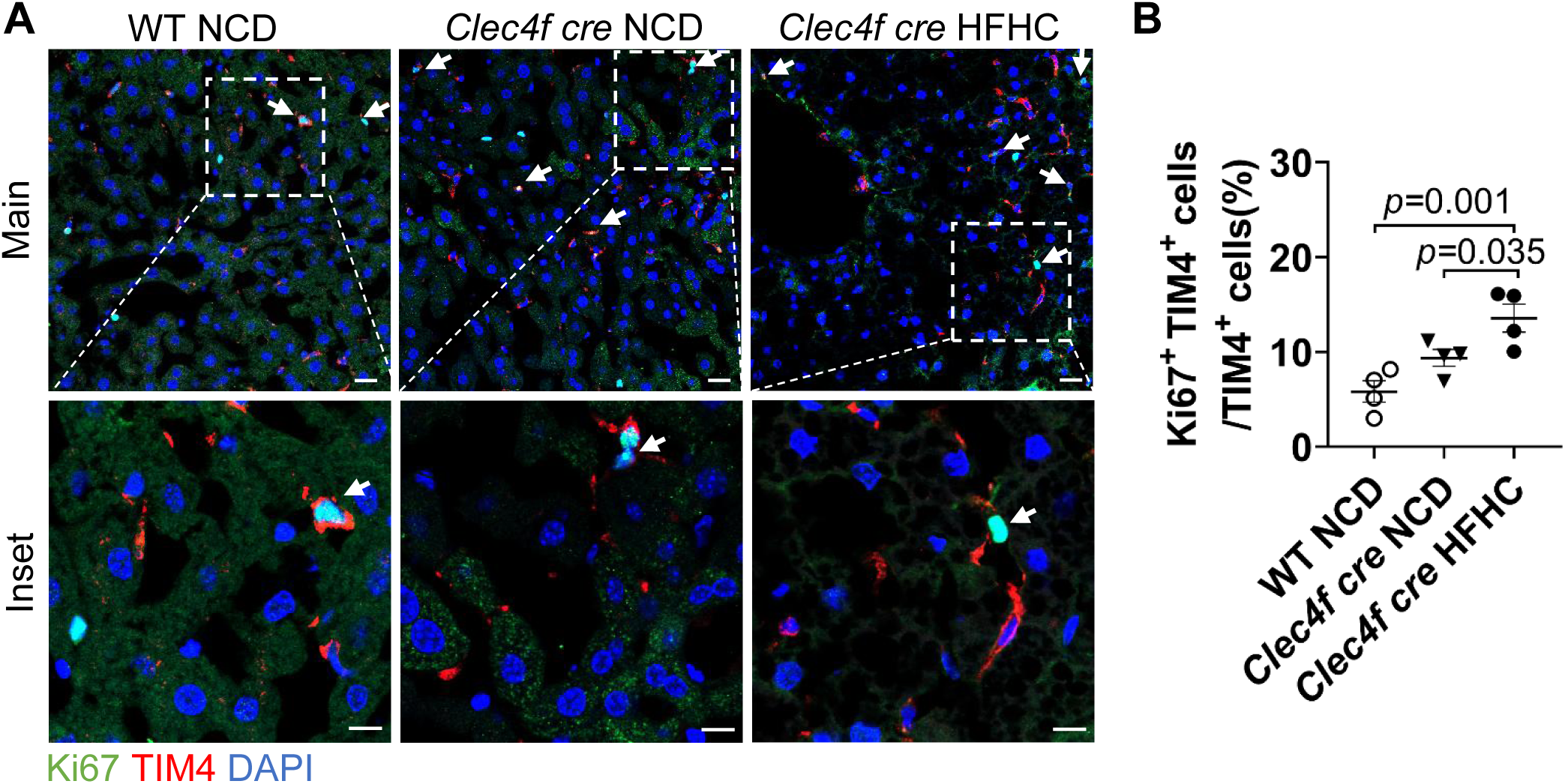
Cre insertion promotes KCs self-proliferation. **(A)** Comparsion of KCs self-proliferation in WT or Clec4f cre mice under NCD or Clec4f cre mice under HFHC diet feeding by costaining of Ki67 (proliferation marker) and TIM4 (KCs marker). Nuclei are counterstained with DAPI (blue). Scale bar: 20μm (main panels) and 10μm (Inset). **(B)** KCs proliferation was quantified. Representative images are shown in A. One-way ANOVA (B). P value as indicated.

