## Supplementary materials and methods for "Hyperactivated Glycolysis Drives Spatially-Patterned Kupffer Cell Depletion in MASLD"

### Isolation of NPCs and KCs

Hepatic nonparenchymal cells (NPCs) were isolated as previously described.1 In brief, the abdominal cavity of mice was promptly opened, and the hepatic portal vein was located after anesthesia. A soft needle was inserted for perfusion with Perfusion buffer (100mL 1× HBSS, 200uL 0.5mol/L EGTA) for 3-5 minutes, while the inferior vena cava was cut to clear the liver. Subsequently, Digestion buffer (60mL 1× HBSS, 120uL 2M $MgSO_4$, 60uL 1.25M $Cacl_2 \cdot H_2O$, 0.015g Collagenase I) was injected at a consistent rate for approximately 15 minutes. Following perfusion, the liver was delicately transferred into a pre-cooled petri dish with PBS, cut into 2-3 mm fragments, and incubated at 37℃ for 15 minutes in Digestion buffer. The liver tissue fluid was then filtered through a 70um cell filter and centrifuged at 50g for 5 minutes (ac/brake=0) at 4℃ to collect the supernatant. This supernatant underwent centrifugation at 450g for 5 minutes (ac/brake=5) at 4℃ to obtain the NPCs population. The NPCs were suspended in a 15mL centrifuge tube with 4mL 20% Optiprep solution, followed by centrifugation at 3000 rpm for 17 minutes (ac/brake=0) at 4℃. After centrifugation, the cells at the boundary between the Optiprep solution and 1xHBSS buffer were collected into a new tube, supplemented with 1xHBSS buffer, and centrifuged for 5 minutes at 450g (ac/brake=5). Red blood cells (RBCs) were lysed with 1mL ACK buffer (150mM $NH_4Cl$, 10mM $KHCO_3$, 0.1mM $Na_2EDTA$, pH=7.2-7.4), neutralized with 1xHBSS buffer, and centrifuged. The cells were resuspended in cell medium and placed in a 3.5cm petri dish for 10 minutes to obtain KCs by washing out un-adherent cells with PBS. The purity and viability of isolated KCs reached 90% and 80%, respectively, which was confirmed by staining with anti-Timd4 antibodies or trypan blue. LDH activity in culture supernatants was measured using a commercial assay kit (Promega, Cat# G1780) according to the manufacturer's instructions. Absorbance was then measured at 490 nm using a microplate reader (Thermo Fisher Scientific, Multiskan Sky).

### KCs Processing and Metabolomic Profiling

The extraction method of cellular metabolites was assessed as previously described[1]. Primary KCs were isolated from livers of HFHC-fed mice at 0, 4, and 8 weeks using established protocols. After a 15-min adherence period, cells were washed twice with ice-cold PBS and detached using 1 mL ice-cold PBS followed by gentle scraping. Cell suspensions were transferred to 1.5 mL tubes, and 3 × $10^6$ cells were pelleted by centrifugation (1,000 × g, 5 min). Supernatants were discarded, and pellets snap-frozen in liquid nitrogen. Metabolites were extracted from frozen pellets with 80% methanol (vol/vol) and analyzed via LC–MS/MS using an AB Sciex 6500 Plus QTRAP mass spectrometer coupled to an ExionLC system. All metabolomic processing and data analysis were performed by BioDeep (https://www.biodeep.cn). Related data are provided in Table S1.

### Flow Cytometry analysis of NPCs

The NPCs were resuspended in fluorescence-activated cell sorting (FACS) buffer, consisting of PBS supplemented with 2% bovine serum albumin. 1uL of anti-CD16/CD32 antibody (Invitrogen, Cat# 14-0161-86) was added to the cell suspension, and the mixture was incubated at 4℃ for 10 min to block any nonspecific binding. After blocking, the Mouse NPCs were labeled with monoclonal antibodies conjugated with fluorescent dyes. This labeling process was carried out at 4℃ for 30 min. The labeled cells were washed thrice with cold FACS buffer. The antibodies used for labeling were as follows: anti-mouse F4/80(APC) (Invitrogen, Cat# 17-4801-82), anti-CD45 (eFluor450) (Invitrogen, Cat# 48-0451-82), anti-mouse Tim-4(PE) (Invitrogen, Cat# 12-5866-82), and anti-mouse CD11b (PerCP/Cyanine5.5) (BioLegend, Cat# 101228). All the primary antibodies were used at a dilution of 1:100. The samples were analyzed by flow cytometry using an LSR Fortessa Cell Analyzer (BD Biosciences). The data obtained were further analyzed using FlowJo version 10.0.

**Preparation of Bone Marrow Derived Macrophages**

To obtain bone marrow-derived macrophages, femur and tibia bone marrow from healthy male *C57BL6/J* was extracted, resuspended in DMEM/F12 medium (VivaCell, Cat# C3113-0500) containing 10% FBS (VivaCell, Cat# C04001-500) and 20% L929 conditioned medium, seeded in culture dishes and cultured at 37 ℃ with 5% $CO_2$ in a humidified atmosphere for 7 days. Fresh medium was added on day 4. The cells were maintained in a standard 37 ℃ in 5% $CO_2$ incubator.

**Isotope Tracing**

$^{13}C$-tracing experiments was assessed as previously described[2]. For $^{13}C$-tracing experiments, BMDMs were isolated from WT and *Chil1*$^{-/-}$ mice, cultured to maturity, and replated. After 12 hours, cells were incubated for 12 hours in glucose-depleted medium supplemented with 10% dialyzed FBS and 15 mmol/L universally labeled [U-$^{13}C$]glucose (Cambridge Isotope Laboratories, CLM-1396-1). Cells were subsequently washed twice with ice-cold glucose-free medium. Following supernatant removal, metabolites were extracted using pre-cooled 80% (vol/vol) methanol for cell lysis.

Metabolite separation was performed using a Vanquish UHPLC system (Thermo Fisher Scientific) equipped with an Amide column (Waters). The mobile phase consisted of: (A) 10 mM ammonium acetate and 0.3% ammonia in water, and (B) 10 mM ammonium acetate and 0.3% ammonia in 90% acetonitrile. Metabolites were separated using a linear gradient elution. Metabolites were ionized, and mass spectrometry data were acquired using an Orbitrap Fusion Tribrid mass spectrometer (Thermo Fisher Scientific). Targeted metabolite analysis was performed by LC–MS/MS using a QTRAP 5500+ system (SCIEX) coupled to an ExionLC AD UHPLC system (SCIEX). Isotope tracing analysis and metabolomic data processing were conducted by BioDeep using the BioDeep Platform (https://www.biodeep.cn). Related data are provided in Table S2.

**Histology & immunofluorescence**

*H&E staining* Tissues were fixed with buffered 10% paraformaldehyde (Sangon Biotech, Cat# A500684-0500) overnight at 4°C and embedded in paraffin. Ultra-thin tissue slices (5μm) were prepared and deparaffinized. H&E staining was performed on the tissue sections, and the slides were examined under a microscope (Olympus, BP80).

*Immunofluorescence on frozen section* Immunofluorescence staining was conducted on frozen sections of fresh liver tissues. Initially, the tissues were fixed using 2% paraformaldehyde for 1 h and subsequently dehydrated overnight in a 30% sucrose solution. The following day, tissue embedding was performed using OCT (SAKURA, Cat# 4583), after which ultra-thin sections of 5μm were sliced. Permeabilization was achieved using 0.02% Triton X-100 for 10 minutes at room temperature, followed by blocking with 5% normal goat serum (VivaCell，Cat# C2530-0100).

*Co-staining of Cleaved caspase 3 with Clec4f* The sections were incubated with primary antibodies including anti-mouse Clec4f (Biolegend, Cat# 156804, 1:300, Alexa Fluor 647) and Cleaved caspase 3 (Cell Signaling, Cat# 9664S, 1:300). The secondary antibodies used were 488-conjugated Affinipure Goat Anti-Rabbit IgG(H+L) (Jackson ImmunoResearch, 111-545-003, 1:600) to lable the Cleaved caspase 3. Nuclei were stained with DAPI (Beyotime, c1006, prediluted), and images were captured using a confocal laser scanning microscope (ZEISS, LSM900).

*Co-staining of TIM4 and F4/80 with Chi3l1* The sections were incubated with primary antibodies including anti-mouse TIM4 (Biolegend，Cat# 130008, 1:300, Alexa Fluor 647), anti-mouse F4/80(Biolegend，Cat# 123140, 1:300, Alexa Fluor 594) and Chi3l1 (Abcam, ab180569, 1:400). The secondary antibodies used were 488-conjugated Affinipure Goat Anti-Rabbit IgG(H+L) (Jackson ImmunoResearch, 111-545-003, 1:600) to lable the Chi3l1. Nuclei were stained with DAPI (Beyotime, c1006, prediluted), and images were captured using a confocal laser scanning microscope (ZEISS, LSM900).

*Co-staining of Clec4f, TIM4, HNF4a, Desmin, IBA1, GS and ki67 with Tunel* Tunel staining was performed on separate frozen sections of liver tissues. Following fixation with 4% paraformaldehyde and treatment with Proteinase K (20 μg/mL in PBS, BBI, Cat# B600169-0002), the sections were permeabilized with Triton X-100 and blocked with normal goat serum. Subsequently, the sections were incubated with an anti-mouse Clec4f antibody (BioLegend，Cat# 156804, 1:300, Alexa Fluor 647), anti-mouse TIM4 (Biolegend，Cat# 130008, 1:300, Alexa Fluor 647), HNF4α(Abcam, Cat# ab181604, 1:400), Desmin(proteintech, Cat#15620-1-AP,1:400), IBA1(Fujifilm, Cat# 019-19741,1:300), GS(Proteintech, Cat#11037-2-AP, 1:400) and Ki67(Abcam, Cat# ab15580, 1:300). The secondary antibodies used were 488-conjugated Affinipure Goat Anti-Rabbit IgG(H+L) (Jackson ImmunoResearch, 111-545-003, 1:600), followed by Tunel staining (servicebio, Cat# G1502-100T) according to the manufacturer's instructions. Nuclei were counterstained with DAPI, and images were captured using a confocal laser scanning microscope (ZEISS, LSM900).

*Immunohistochemical Localization of Chi3l1*

Liver frozen sections from male C57BL/6J mice fed an HFHC diet for 16 weeks were sectioned serially (3μm). Consecutive sections were independently stained using standard immunofluoresence protocols for: Chi3l1 (Abcam, ab180569, 1:400). Lineage markers:

HNF4α (Abcam, Cat# ab181604, 1:400), Desmin (proteintech, Cat#15620-1-AP,1:400), Iba1 (Fujifilm, Cat# 019-19741,1:300). The secondary antibodies used were 488-conjugated Affinipure Goat Anti-Rabbit IgG(H+L) (Jackson ImmunoResearch, 111-545-003, 1:600). Nuclei were counterstained with DAPI, and images were captured using a confocal laser scanning microscope (ZEISS, LSM900). Cellular localization was assigned by aligning morphological features across sequentially stained sections using nuclear and cytoplasmic landmarks.

*Immunofluorescent staining on KCs* KCs were seeded in 24-well plates and subjected to various treatments. In one set of experiments, cells were treated with either DMSO (1μl, Sangon Biotech, Cat# A100231-0500) as a control or oligomycin (1mM in 1μl, Abcam, Cat#ab141829) for 24 h. Another experiment involved treatment with DMSO or PS48 (20 mM in 1μl, Sigma, Cat# P0022) for the same duration. Additionally, BMDM were treated with different glucose concentrations: 5.5 mM glucose alone, 5.5 mM glucose with palmitic acid (800mM in 1μl, Sigma, Cat# P0500), or 10 mM glucose with palmitic acid, all for 24 h. The cell slides were washed with cold PBS and fixed with 4% paraformaldehyde in PBS for 10 min at room temperature. The cells were permeabilized with 0.02% Triton X-100 for 10 min at room temperature. After blocking with 5% normal goat serum, cells were incubated with primarily antibodies anti-Cleaved caspase 3 (Cell Signaling, Cat# 9664S, 1:300) overnight at 4℃. On the second day, after washing with 0.05% PBST, cells were incubating with 488-conjugated Goat Anti-Rabbit IgG (H+L) (Jackson ImmunoResearch, 111-545-003, 1:1000) for 1 h at room temperature. After rinsing with 0.05% PBST, cells were counterstained with DAPI (beyotime, C1006) and mounted onto slides. Images were captured using a confocal laser scanning microscope (ZEISS, LSM900). The identification of KCs follows the above-mentioned experimental protocol. Primary antibodies against TIM4 (Biolegend, Cat# 130008, 1:300, Alexa Fluor 647) are employed to label KCs.

*Calcein-AM staining on KCs* Live cell staining using Calcein-AM was conducted following the manufacturer's instructions from a commercially available kit (Beyotime, Cat# C2015M). KCs were seeded in 12-well plates and subjected to various treatments. In one set of experiments, cells were treated with either DMSO (1μl, Sangon Biotech, Cat# A100231-0500) as a control or oligomycin (1mM in 1μl, Abcam, Cat#ab141829) for 24 h. Another experiment involved treatment with DMSO or PS48 (20 mM in 1μl, Sigma, Cat# P0022) for the same duration. Additionally, BMDM were treated with different glucose concentrations: 5.5 mM glucose alone, 5.5 mM glucose with palmitic acid (800mM in 1μl, Sigma, Cat# P0500), or 10 mM glucose with palmitic acid, all for 24 h.

*Oil Red O staining* Oil red O staining was conducted on unfixed frozen sections embedded directly in OCT. Frozen sections of the liver were cut at a thickness of 10 μm. After rinsing with water, the sections were immersed in 60% isopropanol for 2 min. Subsequently, the sections were stained with an Oil Red O staining solution (Solarbio, Cat# IO1720) at 37°C for 10 to 15 min. Following staining, the sections were immediately placed in 60% isopropanol and washed to 3-5 times to eliminate excess dye solution. Nuclei were counterstained with a hematoxylin staining solution. After rinsing with distilled water, the sections were sealed with glycerol gelatin (Solarbio, Cat# S2150).

*Sirius red staining* Tissues were fixed with buffered 10% paraformaldehyde (Sangon Biotech, Cat# A500684-0500) overnight at 4°C and embedded in paraffin. Ultra-thin tissue slices (5μm) were prepared and deparaffinized. Sirius red staining (Solarbio, Cat#G1472) was performed according to the manufacturer's instructions, and the slides were examined under a microscope (Olympus, BP80).

**Spatial Analysis of KCs Death**

To evaluate the spatial distribution of KCs death along the portal-central axis, the distance between the portal vein (PV) and central vein (CV) was measured in histological sections. The PV-CV axis was systematically divided into three equidistant zones (periportal, intermediate, and pericentral) for regional analysis. KCs death was quantified by isolating fluorescence signals corresponding to cell death markers (TUNEL) through channel thresholding and noise reduction. The positive area and integrated fluorescence intensity were measured within each zone, excluding vascular structures to focus on parenchymal KCs populations. This zonal approach enabled comparative assessment of KCs death patterns across different hepatic microenvironments.

**Diagnosis of MASLD activity score**

Murine MASLD activity was assessed histologically using the NAFLD Activity Score (NAS) on hematoxylin and eosin (H&E) stained liver sections following features: hepatocyte ballooning degeneration (0-2), lobular inflammation (0-3), and steatosis grade (0-3)[3]. The individual scores were summed to yield the total NAS (range 0-8) per animal. A NAS ≥ 5 was considered diagnostic for steatohepatitis (MASH), NAS ≤ 3 indicated not-MASH, and NAS = 4 was indeterminate.

**Extracellular acidification rate (ECAR) measurement**

The ECAR measurements were conducted according to the manufacturer’s instructions (Agilent, Cat# 103020-100). Briefly, BMDM were cultured in a Seahorse XF24 cell culture plate. After 12 h of culture, the DMEM culture medium was pre-treated for 24 h. 1 h before the analysis, the culture medium was changed to the corresponding XF basal medium (Agilent, Cat#103334-100) supplemented with glutamine (The final concentration was 2mmol/L, Agilent, Cat#103579-100), and the culture plates were incubated at 37℃ without $CO_2$. Compounds were added in the following order: 10mM glucose, 1.0 mM oligomycin, and 50mM 2-deoxyglucose. Measurements were conducted using a Seahorse XF24 analyzer (Agilent Technologies). After completing the SeahorseXF glycolytic stress test, basic glycolytic ability and total glycolytic ability were calculated based on the generated report.

**Lactate dehydrogenase (LDH) release assay for cytotoxicity measurement**

The lactate dehydrogenase measurements were conducted according to the manufacturer’s instructions (Promega, Cat# G183A). An appropriate amount of cells were inoculated into the culture plate and treated according to the experimental plan. At the scheduled detection time, 50 μL of supernatant from each well was added to a 96-well plate, and then 50 μL of working solution was added to each well, and incubated at room temperature in the dark for 30 min. Immediately after the reaction, 50μL of termination solution was added to each well to terminate the reaction. Immediately thereafter, the absorbance of the reaction was measured at 490 nm using an Enzyme-labeled instrument (Thermo scientific).

**Western blot analysis**

The sample preparation and procedure were conducted as previously described.[4] The following antibodies were used: anti-Caspase 3 (Cell Signaling Technology, Cat# 9662S, 1:1000), anti-Cleaved Caspase-3 (Cell Signaling Technology, Cat# 9664S, 1:1000), anti-β-actin (Proteintech， Cat# 66009-1-lg, 1:1000). Peroxidase-conjugated Affinipure Goat Anti-Mouse IgG(H+L) (Jackson ImmunoResearch, 115-035-003, 1:2000), and Peroxidase-conjugated Affinipure Goat Anti-Rabbit IgG(H+L) (Jackson ImmunoResearch, 111-035-003, 1:2000) were used for secondary antibody incubation.

**RNA extraction and quantitative real-time PCR**

The TRIzol reagent (Invitrogen, Cat# 15596018) was used to lyse cells or ground tissues, and the resulting mixture was centrifuged at 12,000 rpm/min at 4℃ for 10 min. The liquid portion (supernatant) was carefully transferred to a new tube. To this tube, 200μL of chloroform was added and mixed thoroughly. The tube was left at room temperature for 10 min and subsequently centrifuged. The resulting supernatant was collected in a new tube. Isopropanol, in the same volume as the supernatant, was added to the tube and mixed thoroughly. After incubating for 10 min at 4℃, the mixture was centrifuged. The supernatant was then carefully discarded. RNA was precipitated by adding 75% ethanol to the tube, followed by centrifugation. This process was repeated. The tube was dried for 10 min at room temperature, and an appropriate amount of ribonuclease-free water was added to dissolve the RNA precipitate. The concentration and purity of RNA samples were determined using an ultraviolet spectrophotometer (NanoDrop). The RNA template was subjected to reverse transcription using a cDNA first strand synthesis kit (Takara, Cat# 6210B). Quantitative PCR was performed using SYBR Green Master Mix (Thermo Fisher, Cat# A25742) in triplicates following the manufacturer's instructions. This was performed on a Real-Time PCR QuatStudio1 with accompanying software, following the instructions provided by the manufacturer (Life Technologies, Grand Island, NY, USA). Primer sequences are provided in Table S3.

**Table S3**

**qPCR primers**

| Targeted gene | Primer | Sequence |
|---|---|---|
| musSlc2a1 | F(5‘-3’) | CAGTTCGGCTATAACACTGGTG |
| | R(5‘-3’) | GCCCCCGACAGAGAAGATG |
| musHk3 | F(5‘-3’) | CTGAGTCAAGGCTGTATCCTCC |
| | R(5‘-3’) | TGCACCAGTTCAGCATCTGAGG |

| | | |
|---|---|---|
| musPfkfb3 | F(5‘-3’) | CCCAGAGCCGGGTACAGAA |
| | R(5‘-3’) | GGGGAGTTGGTCAGCTTCG |
| musPkm | F(5‘-3’) | GCCGCCTGGACATTGACTC |
| | R(5‘-3’) | CCATGAGAGAAATTCAGCCGAG |
| mus6pgd | F(5‘-3’) | CATCGCTGCAAAAGTGGGAACC |
| | R(5‘-3’) | AGCCTCACAGATGAGCTGCATG |
| musG6pd | F(5‘-3’) | GACCAAGAAGCCTGGCATGTTC |
| | R(5‘-3’) | AGACATCCAGGATGAGGCGTTC |
| musPygl | F(5‘-3’) | GGCAGAAGTGGTGAACAATGACC |
| | R(5‘-3’) | TCCGATAGGTCTGTGGCTGGAA |
| musUgp2 | F(5‘-3’) | CTGATGAACCCACCCAATGGGA |
| | R(5‘-3’) | GAGCGATTTCCACCAGTCTCAG |
| musGys1 | F(5‘-3’) | CACAGAACGGTTGTCGGACTTG |
| | R(5‘-3’) | AGGTGAAGTGGTCTGGAAAGGC |
| musIdh1 | F(5‘-3’) | CAGGCTCATAGATGACATGGTGG |
| | R(5‘-3’) | CACTGGTCATCATGCCAAGGGA |
| musOgdh | F(5‘-3’) | GGTGTCGTCAATCAGCCTGAGT |
| | R(5‘-3’) | ATCCAGCCAGTGCTTGATGTGC |
| musAcadm | F(5‘-3’) | AGGGTTTAGTTTTGAGTTGACGG |
| | R(5‘-3’) | CCCCGCTTTTGTCATATTCCG |
| musHadh | F(5‘-3’) | TTCCAGAGGCTGGACAAGTTCG |
| | R(5‘-3’) | GCCAGCAAATCGGTCTTGTCTG |
| musNos2 | F(5‘-3’) | GAGACAGGGAAGTCTGAAGCAC |
| | R(5‘-3’) | CCAGCAGTAGTTGCTCCTCTTC |
| musCxcl9 | F(5‘-3’) | CCTAGTGATAAGGAATGCACGATG |
| | R(5‘-3’) | CTAGGCAGGTTTGATCTCCGTTC |
| musCIITA | F(5‘-3’) | ACCTTCGTCAGACTGGCGTTGA |
| | R(5‘-3’) | GCCATTGTATCACTCAAGGAGGC |
| MusCD86 | F(5‘-3’) | ACGTATTGGAAGGAGATTACAGCT |
| | R(5‘-3’) | TCTGTCAGCGTTACTATCCCGC |
| musCcl3 | F(5‘-3’) | ACTGCCTGCTGCTTCTCCTACA |
| | R(5‘-3’) | ATGACACCTGGCTGGGAGCAAA |
| musCcl5 | F(5‘-3’) | CCTGCTGCTTTGCCTACCTCTC |
| | R(5‘-3’) | ACACACTTGGCGGTTCCTTCGA |
| musChil3 | F(5‘-3’) | CTCCAGTGTAGCCATCCTTAGG |
| | R(5‘-3’) | TACTCACTTCCACAGGAGCAGG |
| musRetnla | F(5‘-3’) | CCAAGATCCACAGGCAAAGCCA |
| | R(5‘-3’) | CAAGGAACTTCTTGCCAATCCAG |
| musArg1 | F(5‘-3’) | GCTGAAGGTCTCTTCCATCACC |
| | R(5‘-3’) | CATTGGCTTGCGAGACGTAGAC |
| musMrc1 | F(5‘-3’) | GTTCACCTGGAGTGATGGTTCTC |
| | R(5‘-3’) | AGGACATGCCAGGGTCACCTTT |
